# Golgi retention and oncogenic KIT signaling via PLCγ2-PKD2-PI4KIIIβ activation in GIST cells

**DOI:** 10.1101/2022.12.19.520889

**Authors:** Yuuki Obata, Kazuo Kurokawa, Takuro Tojima, Miyuki Natsume, Isamu Shiina, Tsuyoshi Takahashi, Ryo Abe, Akihiko Nakano, Toshirou Nishida

**Author notes:** Corresponding author: Yuuki Obata, Ph.D., Laboratory of Intracellular Traffic & Oncology, National Cancer Center Research Institute, Tsukiji, 5-1-1, Chuo-ku, Tokyo, 104-0045, Japan. **E-mail**; Kazuo Kurokawa; Takuro Tojima; Miyuki Natsume; Isamu Shiina; Tsuyoshi Takahashi; Ryo Abe; Akihiko Nakano; Toshirou Nishida.

## Abstract

Most gastrointestinal stromal tumors (GISTs) develop due to gain-of-function mutations in the tyrosine kinase, *KIT*. We recently showed that mutant KIT mislocalizes to the Golgi area and initiates uncontrolled signaling. However, the molecular mechanisms underlying its Golgi retention remain unknown. Here, we show that protein kinase D2 (PKD2) is activated by the mutant, which causes KIT’s Golgi retention. In PKD2-inhibited cells, KIT migrates from the Golgi region to lysosomes and subsequently undergoes degradation. Importantly, delocalized KIT is unable to trigger downstream activation. In the Golgi area, KIT activates the PKD2-phosphatidylinositol 4-kinaseIIIβ (PKD2-PI4KIIIβ) pathway through phospholipase γ2 (PLCγ2) to generate a PI4P-rich membrane domain, where the AP1-GGA1 complex is aberrantly recruited. Disruption of any factors in this cascade results in KIT release from the Golgi region, indicating that these PKD2-related pathways are responsible for the Golgi retention of KIT. Our findings unveil the molecular mechanisms underlying KIT mislocalization and provide evidence for a new strategy for inhibition of oncogenic signaling.

## INTRODUCTION

KIT, platelet-derived growth factor receptor A/B (PDGFRA/B), and FMS-like tyrosine kinase 3 (FLT3) are class III receptor tyrosine kinases (RTKs)^1^. Upon ligand stimulation at the plasma membrane (PM), they initiate tyrosine phosphorylation signaling, leading to cell proliferation and survival in hematopoietic cells, mast cells, and interstitial cells of Cajal^2, 3^.

KIT is composed of an amino-terminal extracellular domain, a transmembrane domain, a juxta-membrane region, and a carboxy-terminal cytoplasmic kinase domain^4^. Upon ligand binding, KIT autophosphorylates specific tyrosine residues, such as Y703, resulting in the recruitment of downstream molecules^5^. Phospho-KIT activates the PI3-kinase-AKT (PI3K-AKT) pathway and signal transducer and activator of transcription proteins (STATs), which control protein expression and metabolism^4^. Thus, gain-of-function mutations in *KIT* lead host cells to undergo uncontrolled growth, resulting in the development of acute myelogenous leukemia (AML), mast cell leukemia (MCL), and gastrointestinal stromal tumor (GIST)^6–9^. In particular, gene alterations in *KIT* in the kinase domain (*exon 17* (*ex17*): *D816V,* etc.) or juxta-membrane region (*ex11*: *deletion*, etc.), which cause constitutive kinase activation, are found in most cases of MCL and more than 70% of GIST, respectively^9,10^. A tyrosine kinase inhibitor (TKI), imatinib (IMA), dramatically improved the prognosis of patients with advanced GIST bearing *KIT ex11* mutations^11^. However, secondary mutations in *KIT ex13* (*V654A*) or *ex17*, which lose sensitivity to TKI, occur in IMA-treated GIST patients within 2–3 years^11, 12^. Thus, further understanding of RTK signaling is required for the development of effective targeted therapies.

We previously reported that in MCL and AML, KIT bearing ex11 or ex17 mutations aberrantly localizes to endosome-lysosomal compartments^13–15^. In sharp contrast, during biosynthetic trafficking in GIST cells, the KIT^Ex17^ mutant and KIT^Ex11^ accumulate in the Golgi/*trans-*Golgi network (TGN) area, where they initiate oncogenic signaling^16–18^. We further showed that FLT3-internal tandem duplication (FLT3-ITD) in AML triggers growth signals in the Golgi area and the endoplasmic reticulum (ER)^19^. Although the organelles where RTK accumulates differ among cancer types, compartment-dependent signaling is a characteristic feature of mutant RTKs. However, the molecular mechanism underlying RTK organelle retention remains unclear.

In this study, we show that protein kinase D2 (PKD2, a membrane fission regulator^20–22^) and its effectors trap KIT mutants in the Golgi area of GIST cells. Among the multiple compounds tested, only the PKD inhibitor releases KIT from the Golgi region, indicating that PKD arrests KIT in the Golgi/TGN. After release from the Golgi region, KIT migrates to lysosomes via the PM and rapidly undergoes degradation. Importantly, KIT that is delocalized from the Golgi region is unable to activate downstream molecules in both IMA-sensitive and IMA-resistant GIST cells. Among PKD1–3, only PKD2 plays a critical role in KIT retention, and it is activated and associated with KIT in the Golgi area. KIT requires PLCγ2 to activate PKD2. Furthermore, PI4KIIIβ is an effector of KIT-PLCγ2-PKD2, and it generates PI4P for recruitment of adaptor protein complexes (APs) and Golgi-associated, γ-adaptin ear-containing, ARF-binding proteins (GGAs). Interestingly, knockdown of PLCγ2, PI4KIIIβ, γ-adaptin (AP1 component), or GGA1 phenocopies PKD2 knockdown in the release of KIT from the Golgi region. Thus, these aberrantly recruited factors trap KIT in the Golgi area. In contrast, these pathways are not involved in RTK organelle retention in leukemia cells. Therefore, our study demonstrates that the PLCγ2-PKD2-PI4KIIIβ-AP1-GGA1 cascade plays a critical role in Golgi retention of KIT in GISTs. Our findings uncover one of the molecular mechanisms underlying oncogenic mutant KIT organelle retention and provide evidence of a new strategy for the inhibition of oncogenic signaling.

## RESULTS

### KIT is retained in the Golgi area in a PKD activity-dependent manner in GIST cells

To determine the cause of KIT retention in the Golgi area, we treated GIST-T1 cells (*KIT^Ex11^*, IMA-sensitive), which were established from a GIST patient^23^, with more than 20 compounds followed by immunofluorescence confocal microscopic analyses. As shown in Figure S1A, most compounds had no effect on KIT localization in the Golgi area, similarly to our previous report that PP2 (SRC tyrosine kinase inhibitor) and AKT inhibitor VIII did not affect the mutant localization^16^. Interestingly, upon 8-h treatment with CRT0066101 (CRT), a PKD inhibitor, KIT migrated from the Golgi region to non-Golgi punctate structures (Figures 1A and S1A), as confirmed by golgin97 or Golgi matrix protein 130 kDa (GM130) staining (Figures 1B and S1B). After 16-h CRT treatment, KIT was barely detected in the Golgi area (Figures 1A, 1B, and S1C). Immunoblotting revealed that the inhibitor decreased KIT protein levels in a time-dependent manner without affecting mutant autophosphorylation (Figure 1C). Inhibition of PKD activity was confirmed by measuring the decrease in phospho-PKD substrate levels (Figure 1C, bottom panels). Importantly, because KIT in CRT-treated cells moved out from the Golgi region, the mutant signaling site, the activation of AKT and STAT5 also went down (Figure 1C). We next hypothesized that CRT treatment caused translocation of KIT to lysosomes. Indeed, KIT was found in lysosomal-associated membrane protein 1-positive (LAMP1-positive) lysosomes in PKD-inhibited cells (Figure 1D). Phospho-KIT^Y703^ (pKIT^Y703^), which mainly represents the mutant form, was also located in lysosomes of CRT-treated cells (Figures 1E and S1C). Furthermore, the decrease of KIT level was suppressed by NH4Cl or chloroquine, which blocks lysosomal protein degradation (Figure 1F). In contrast, in the presence of CRT, the two lysosomal inhibitors did not affect the levels of the Golgi/TGN proteins, GM130, golgin97, and syntaxin 6 (Figure S1D), supporting that PKD inhibition selectively causes migration of KIT to lysosomes and subsequent degradation. Next, we investigated whether similar results were obtained from IMA-resistant GIST cells (GIST-R9, *KIT^Ex11/Ex17^*; GIST430, *KIT^Ex11/Ex13^*; GIST48, *KIT^Ex11/Ex17^*) (refs.^24, 25^). Like GIST-T1 KIT, IMA-resistant mutants were localized and autophosphorylated in the Golgi area (Figure S1E; refs. ^17, 18^). IMA-resistant KIT level was decreased by CRT treatment (Figures 1G and S1F), resulting in AKT and STAT5 inactivation. PKD was proposed to play a role in generation of membrane carriers from the TGN^20–22, 26, 27^. Considering that PKD inhibition releases KIT mutant from the Golgi region, we hypothesized that aberrant PKD activation causes excessive recruitment of the components that are involved in membrane fission in the Golgi/TGN area, resulting in KIT being trapped there.

**Figure 1.**
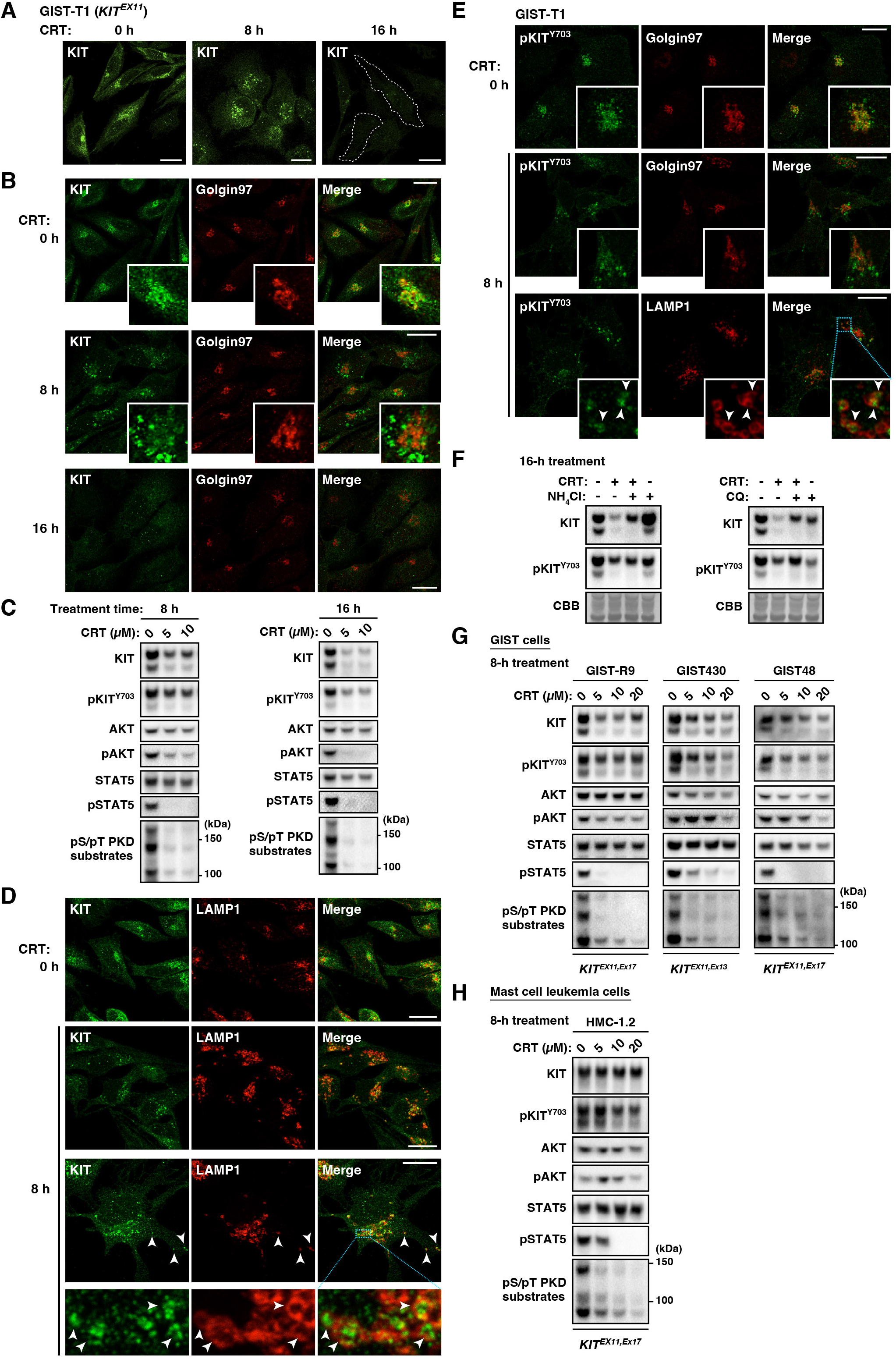
CRT0066101, a PKD inhibitor, rescues Golgi retention of mutant KIT and moves the mutant to lysosomes in GIST cells. (A–E) GIST-T1 cells were treated with 10 µM CRT0066101 (CRT, PKD inhibitor) for the indicated periods. (A and B) Cells were immunostained for KIT (green) and golgin97 (red). Insets show magnified images of the Golgi area. Dashed lines indicate cell borders. Bars, 20 µm. NB: KIT migrates from the Golgi region to non-Golgi punctate structures in the presence of CRT. (C) Lysates were immunoblotted for KIT, phospho-KIT^Y703^ (pKIT^Y703^), AKT, pAKT, pSTAT5, and phospho-PKD substrates (pS/pT PKD substrates). (D) Cells were immunostained for KIT (green) and LAMP1 (lysosomal-associated membrane protein 1, red). Magnified images of the boxed area are shown. Arrowheads indicate KIT in lysosomes. Bars, 20 µm. NB: KIT was found inside LAMP1-positive lysosomes when PKD was inhibited. (E) Cells were immunostained for pKIT^Y703^ in conjunction with golgin97 or LAMP1. Magnified images of the boxed area are shown. Arrowheads indicate pKIT^Y703^ in lysosomes. Bars, 20 µm. (F) GIST-T1 cells were treated with 10 µM CRT and/or 20 mM NH_4_Cl, 100 µM chloroquine (CQ) for 16 hours, and immunoblotted. Total protein levels were confirmed by Coomassie Brilliant Blue (CBB) staining. (G and H) Imatinib-resistant GIST cells (G) or HMC-1.2 cells (H) were treated with CRT for 8 hours, and immunoblotted.

We previously showed that mutant KIT in MCL passes through the Golgi/TGN and accumulates on endosome-lysosome membranes^13–15^. As shown in Figure 1H, unlike KIT in GIST cells, KIT was not degraded by CRT treatment in the MCL cell line, HMC-1.2. Phospho-STAT5 expression was diminished by the treatment, but this was not due to KIT downregulation. Taken together, these results suggest that PKD activity is essential for Golgi KIT retention in GIST cells.

### In PKD-inhibited GIST cells, KIT migrates to the PM from the Golgi region, and undergoes EIPA-sensitive endocytosis

Next, we investigated whether the tyrosine kinase activity of KIT is required for the mutant trafficking to lysosomes in PKD-inhibited cells. We previously showed that TKIs (IMA or PKC412) decrease the KIT level in the Golgi/TGN and increase it in the PM^15, 16^ (see also Figure 2A). When GIST-T1 cells were treated with CRT plus IMA for 16 hours, KIT was primarily found on the PM (Figure 2B), suggesting that, in the presence of CRT, KIT migrates to lysosomes via the PM in a kinase activity-dependent manner. We then investigated the mode of endocytosis required for KIT internalization. Intriguingly, in the cells treated with CRT plus 5-[N-ethyl-N-isopropyl] amiloride (EIPA), which blocks endocytosis via sodium-hydrogen exchanger (NHE) inhibition^28^, KIT was found on the PM (Figure 2C). Other endocytosis inhibitors, such as dynasore (dynamin inhibitor) and cytochalasin D (inhibitor of actin polymerization), did not affect KIT localization in the presence or absence of CRT (Figure S2). In CRT-treated cells, the inhibition of endocytosis with IMA or EIPA blocked KIT degradation (Figures 2D and 2E). EIPA is a well-known inhibitor of macropinocytosis, which requires PI3K^29^. To confirm whether KIT endocytosis was macropinocytosis, we treated cells with CRT plus LY294002, a PI3K inhibitor, but did not observe any effect on KIT distribution in the presence or absence of CRT (Figures 2F and S2). These results indicate that KIT undergoes EIPA-sensitive, but LY294002/cytochalasin D/dynasore-insensitive endocytosis. Taken together, we conclude that after release from the Golgi/TGN by PKD inhibition, KIT migrates to the PM and undergoes tyrosine kinase activity-and NHE-dependent endocytosis toward lysosomes.

**Figure 2.**
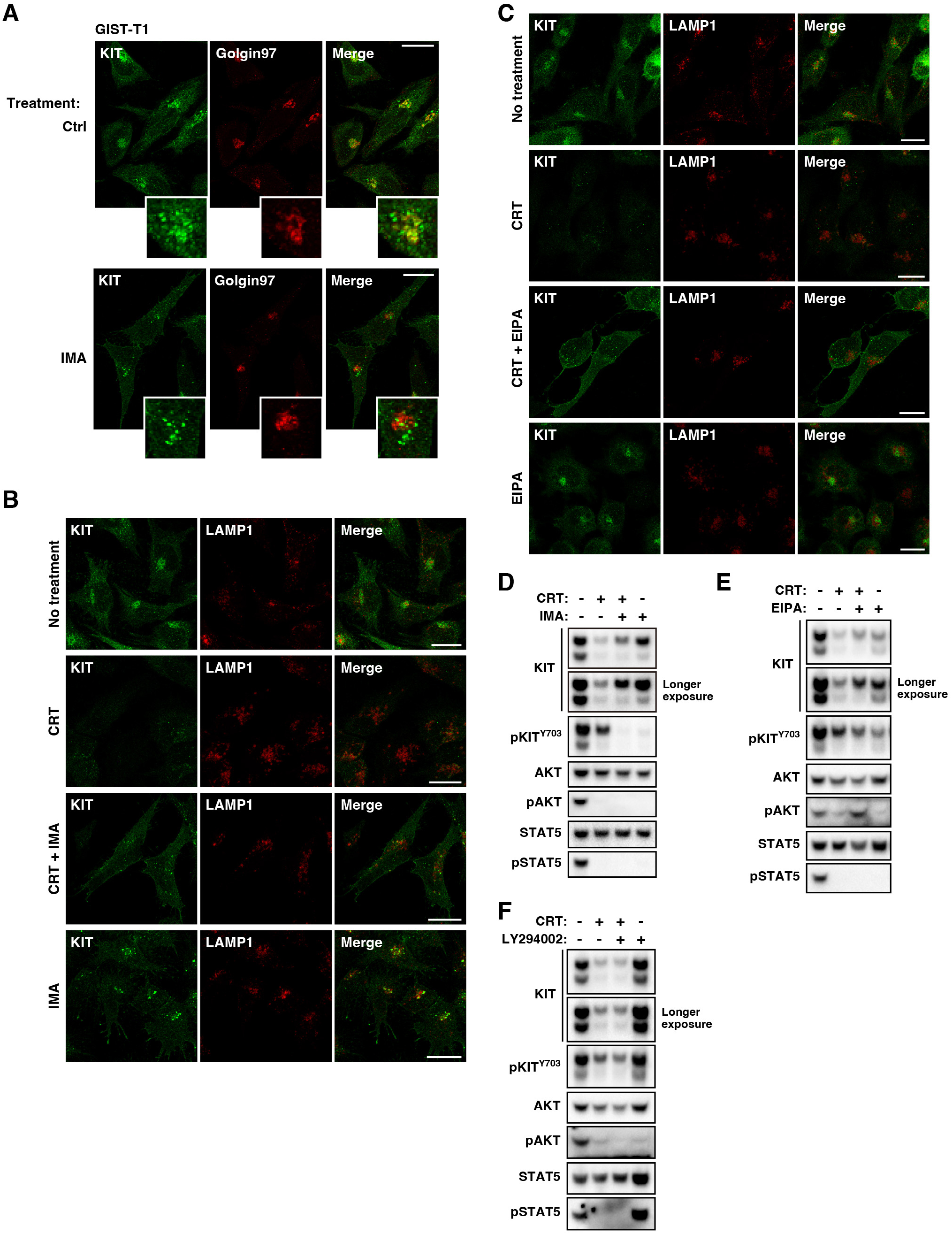
KIT is trafficked to lysosomes via the PM by EIPA-sensitive endocytosis. (A) GIST-T1 cells were treated with 200 nM imatinib (IMA, KIT kinase inhibitor) for 8 hours, then immunostained for KIT (green) and golgin97 (red). Insets show magnified images of the Golgi region. Bars, 20 µm. (B–F) GIST-T1 cells were treated with (B and D) 10 µM CRT0066101 (CRT, PKD inhibitor) and/or 200 nM IMA, (C and E) 40 µM ethyl-isopropyl amiloride (EIPA, inhibitor of Na^+^/H^+^ exchanger), (F) 20 µM LY294002 (PI3-kinase inhibitor) for 16 hours. (B and C) Cells were immunostained for KIT (green) and LAMP1 (lysosome marker, red). Bars, 20 µm. (D–F) Lysates were immunoblotted for KIT, phospho-KIT^Y703^ (pKIT^Y703^), AKT, pAKT, STAT5, and pSTAT5. NB: KIT incorporation into lysosomes induced with CRT treatment was inhibited by EIPA.

### PKD2, but not PKD1/PKD3, plays a critical role in Golgi retention of KIT in GIST cells

Human PKD consists of three members, PKD1, PKD2, and PKD3 (refs. ^30–32^). In GIST-T1 cells, all PKD members were expressed and found in the Golgi area (Figures 3A and S3A). Thus, we performed a small interfering RNA-mediated (siRNA-mediated) knockdown to determine the PKD member critical for KIT retention. Interestingly, PKD2 depletion had similar effects as CRT treatment, but neither PKD1 knockdown nor PKD3 knockdown affected KIT levels (Figure 3B). Moreover, immunofluorescence revealed that KIT was found in lysosomes, but not in the Golgi area in PKD2 knockdown cells (Figure 3C). The movement of KIT to lysosomes is an indication of the delocalization of the mutant from the Golgi region. Figure 3D shows that KIT localized in the Golgi area together with PKD2. Furthermore, IMA-resistant GIST cells showed similar results (Figures 3E, 3F, S3C, and S3D).

**Figure 3.**
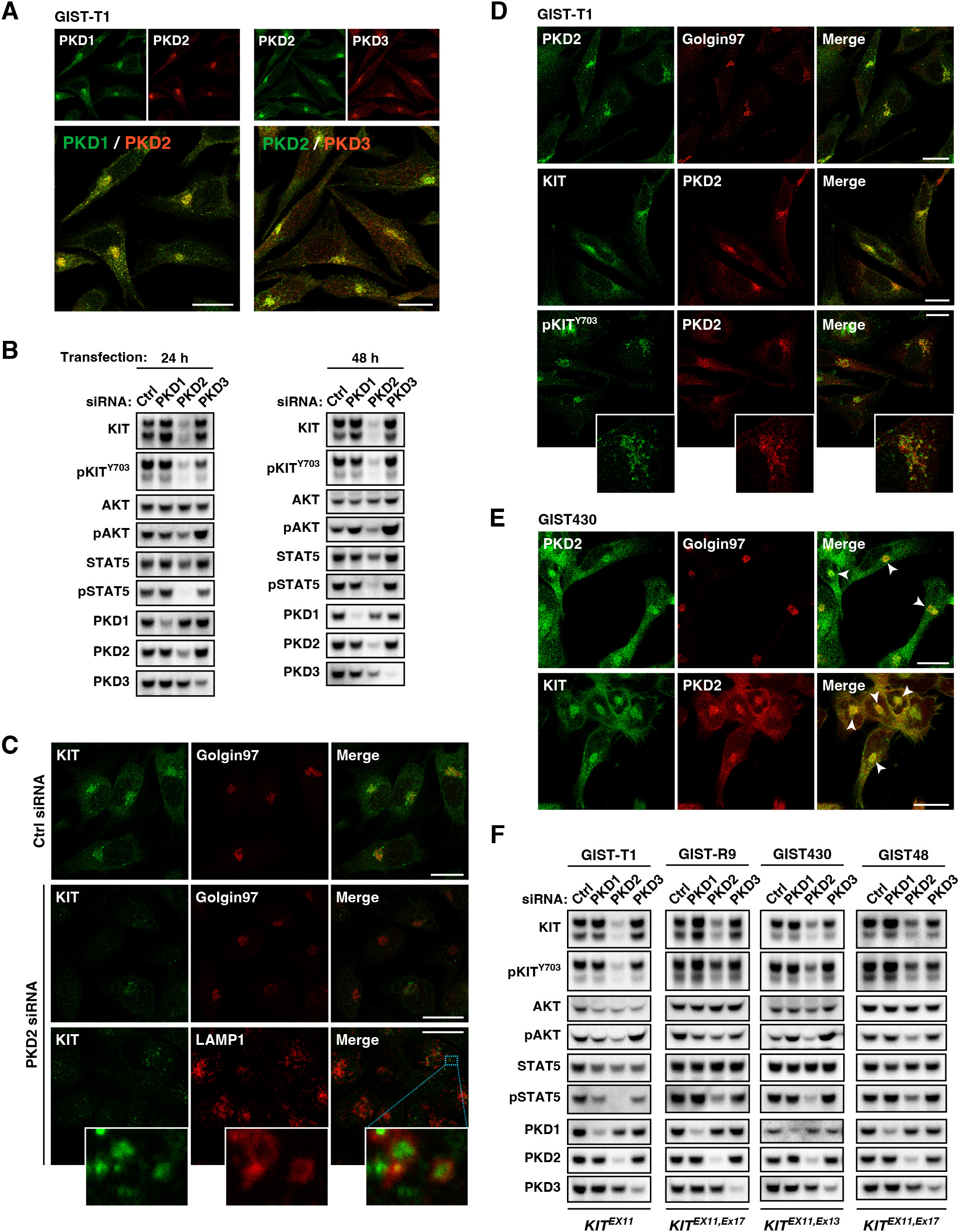
KIT is retained in the Golgi area in a PKD2-dependent manner in GIST cells. (A) GIST-T1 cells were immunostained for PKD1 (green), PKD2 (red or green), or PKD3 (red). Bars, 20 µm. (B) GIST-T1 cells were transfected with the indicated siRNAs for the indicated periods, then immunoblotted for KIT, phospho-KIT^Y703^ (pKIT^Y703^), AKT, pAKT, STAT5, pSTAT5, and PKD1/2/3. NB: The effect of knockdown of PKD2 but not of PKD1/3 on KIT levels was similar to that of CRT0066101 treatment. (C) GIST-T1 cells were transfected with PKD2 siRNA for 30 hours, then immunostained for KIT (green) in conjunction with golgin97 (red) or LAMP1 (lysosome marker, red). Bars, 20 µm. Magnified images of the boxed area are shown. (D and E) GIST-T1 (D) or GIST430 cells (E) were immunostained with the indicated antibodies. Bars, 20 µm. (F) GIST-T1, GIST-R9, GIST430, or GIST48 cells were transfected with the indicated siRNAs for 30 hours, and immunoblotted.

Next, we analyzed the activation of PKD by immunostaining for phospho-PKD2^S876^ (pPKD2^S876^), a marker of the kinase activation^31, 32^. As shown in Figure 4A, pPKD2^S876^ was mainly found in the Golgi area, where KIT is localized. PKD2 phosphorylation in GIST-R9 and GIST430 cells was observed in the Golgi area, similarly to GIST-T1 (Figure S4A). We were unable to detect pPKD2^S876^ in the Golgi area in GIST48 cells by our immunofluorescence, probably due to low phosphorylation levels.

**Figure 4.**
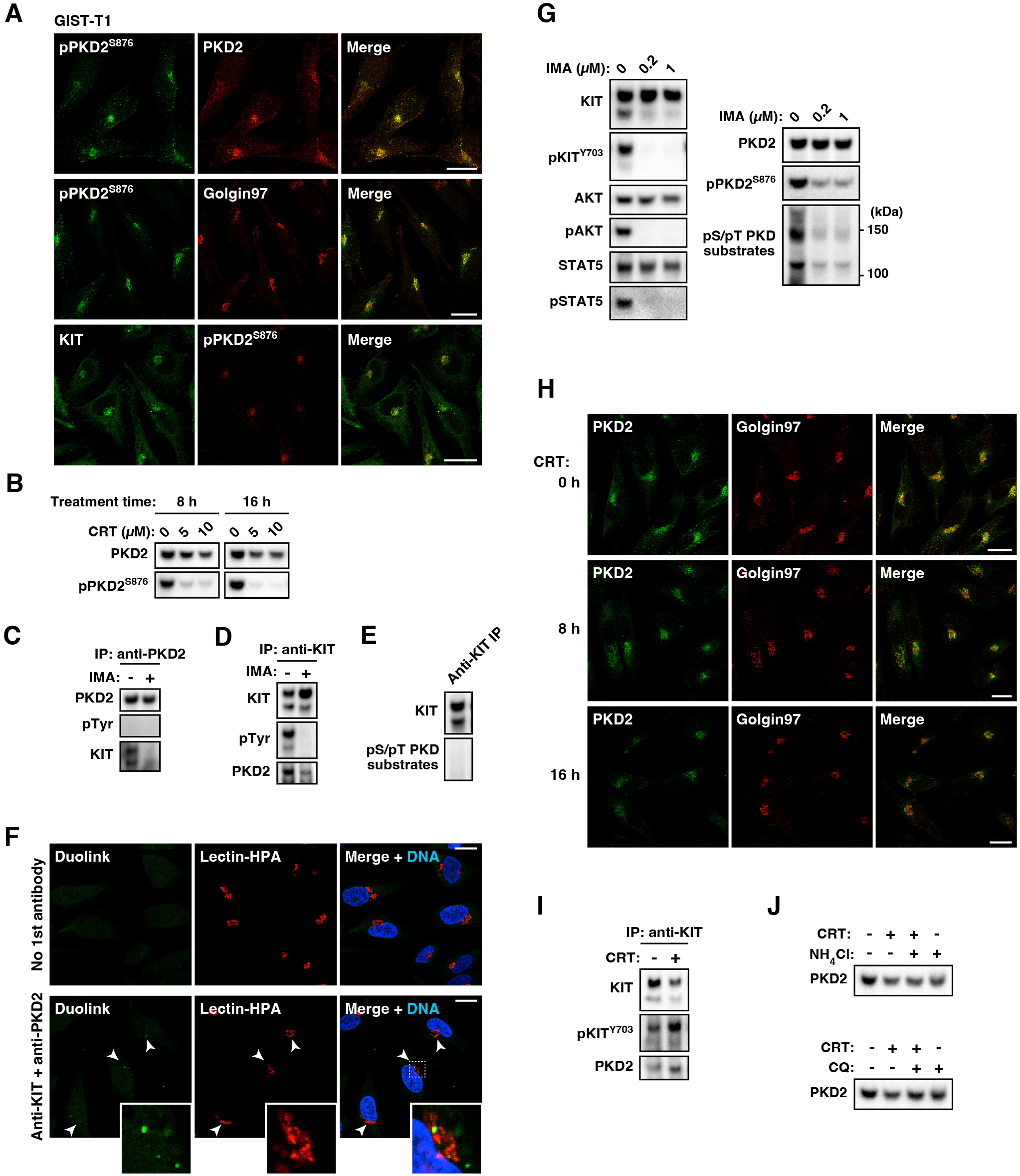
KIT activates PKD2 in the Golgi area in GIST cells. (A) GIST-T1 cells were immunostained for phospho-PKD2^S876^ (pPKD2^S876^, red or green) in conjunction with PKD2 (red), golgin97 (red), or KIT (green). Bars, 20 µm. (B) GIST-T1 cells were treated with CRT0066101 (CRT, PKD inhibitor) for 8 hours (left) or 16 hours (right), then immunoblotted. (C and D) GIST-T1 cells were treated with 200 nM imatinib (IMA, KIT kinase inhibitor) for 8 hours. PKD2 (C) or KIT (D) was immunoprecipitated, and the immunoprecipitates (IPs) were immunoblotted. pTyr, phospho-tyrosine. (E) KIT IPs were immunoblotted for KIT and phospho-PKD substrates. (F) Proximity ligation assay (PLA) was performed with anti-KIT and anti-PKD2 antibodies. Distribution of Duolink PLA signals (green), Lectin-HPA (Golgi marker, red), and nucleus (DAPI, blue) are shown. Insets show magnified images of the boxed area. Bars, 20 µm. (G) GIST-T1 cells were treated with IMA for 8 hours, and immunoblotted. NB: IMA blocked PKD2 activation, resulting in a decrease of phospho-PKD substrates. (H) GIST-T1 cells were treated with 10 µM CRT for the indicated periods, then immunostained for PKD2 (green) and golgin97 (red). Bars, 20 µm. (I) GIST-T1 cells were treated with 10 µM CRT for 8 hours, and KIT was immunoprecipitated with anti-KIT antibody and IPs were immunoblotted. (J) GIST-T1 cells were treated with 10 µM CRT and/or 20 mM NH_4_Cl, 100 µM chloroquine (CQ) for 16 hours, and immunoblotted. Total protein levels were confirmed by Coomassie Brilliant Blue staining in the main Figure 1F.

Commercially available anti-pPKD1^S910^ antibody has been shown to cross-react with pPKD2^S876^ (ref. ^33^; Figure S4B). Our immunoblotting data for GIST-T1 showed that the position of the pPKD1^S910^ band was consistent with that of PKD2 rather than PKD1 (Figure S4C). Considering that only PKD2 knockdown affected KIT levels, PKD2 is likely the major activated PKD member in GIST-T1. Moreover, anti-pPKD1^S910^ antibody stained the Golgi area (Figure S4D). Phospho-PKD2 levels were decreased by CRT treatment (Figures 4B and S4E). Taken together, the effect of CRT on KIT was mainly due to PKD2 inhibition in GIST-T1 cells.

Next, we examined the association between KIT and PKD2 using a co-immunoprecipitation assay. KIT was co-immunoprecipitated with PKD2, and KIT inhibition by IMA blocked this association (Figures 4C and 4D). These results suggest that KIT binds to PKD2 in a kinase activity-dependent manner. We were unable to detect tyrosine phosphorylation in PKD2 (Figure 4C) and KIT was probably not phosphorylated by PKD (Figure 4E), suggesting that they do not directly phosphorylate each other. To determine whether KIT is associated with PKD2 in the Golgi area, we performed a proximity ligation assay (PLA). When anti-KIT and anti-PKD2 antibodies were added, Duolink PLA signals were detected in the perinuclear region, which was stained with a Golgi marker lectin-HPA (Figure 4F). In addition, KIT inhibition with IMA decreased PKD2 activity and phospho-PKD substrate levels (Figures 4G and S4F). Collectively, these results suggest that KIT induces PKD2 activation in the Golgi area.

As shown in Figures 4H, S4G, and S4H, neither CRT nor IMA affected the localization of PKD2 to the Golgi area. Furthermore, PKD2 was not dissociated from KIT by CRT (Figure 4I), indicating that its association with KIT is activity-independent. In the presence of CRT, lysosomal enzyme inhibitors did not change the PKD2 level (Figure 4J), indicating that, unlike KIT, PKD2 localizes to and stays in the Golgi area, regardless of its activation states.

### KIT activates PKD2 via PLCγ2 for its Golgi retention in GIST cells

PKD requires diacylglycerol (DAG) in the Golgi/TGN membrane for its function^21, 26, 34^. Thus, we investigated whether RTK-associated DAG-producing enzymes such as PLCγ1/2 (refs. ^35, 36^), play a role in PKD2 activation. First, we knocked down PLCγ1 or PLCγ2 in GIST-T1 cells followed by immunoblotting. Interestingly, PLCγ2 knockdown showed similar results to PKD2 knockdown (Figure 5A). Although PLCγ2 knockdown did not affect pAKT in GIST430 cells, KIT and pSTAT5 levels were decreased (Figure S5A). Importantly, PLCγ2 knockdown decreased pPKD2^S876^ and the intracellular levels of phospho-PKD substrates in GIST-T1 and GIST430 cells (Figures 5A and S5A), indicating that PLCγ2 lies upstream of PKD2. As shown in Figure 5B, KIT looked as if engulfed in lysosomes in PLCγ2-knocked down GIST-T1 cells. Phospho-PLCγ2^Y759^, a marker of the lipase activation^35^, was predominantly located in the Golgi area (Figure 5C), although PLCγ2 protein was not detected by our immunofluorescence. KIT kinase inhibition by IMA decreased pPLCγ2^Y759^ expression (Figure 5D), indicating that PLCγ2 mediates the interaction between KIT and PKD2. Furthermore, in the proximity ligation assay, Duolink PLA signals of KIT-pPLCγ2^Y759^ were observed in the Golgi area, which were diminished by IMA treatment (Figure 5E). Collectively, these results suggest that KIT activates PKD2 through PLCγ2 and thus causes Golgi retention in GIST cells.

**Figure 5.**
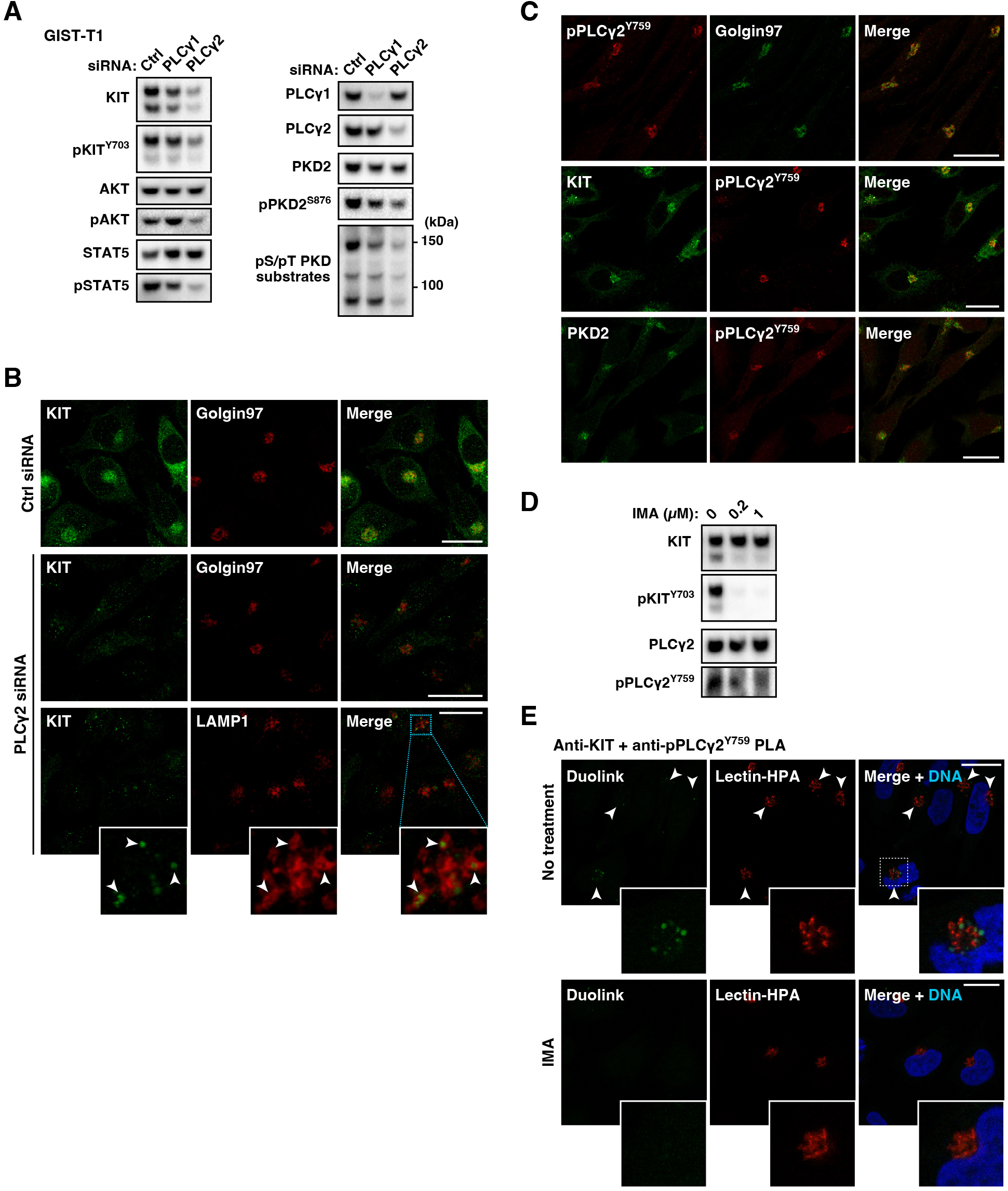
KIT activates PKD2 via PLCγ2 in the Golgi area in GISTs cells. (A and B) GIST-T1 cells were transfected with the indicated siRNAs for 30 hours. (A) Lysates were immunoblotted. (B) Cells were immunostained for KIT (green) in conjunction with golgin97 (red) or LAMP1 (lysosome marker, red). Magnified images of the boxed area are shown. Arrowheads indicate KIT in lysosomes. Bars, 20 µm. NB: Knockdown of PLCγ2 phenocopied that of PKD2 in KIT reduction and signal inhibition. (C) GIST-T1 cells were immunostained for phospho-PLCγ2^Y759^ (pPLCγ2^Y759^, green or red) in conjunction with golgin97 (red), KIT (green), or PKD2 (green). Bars, 20 µm. (D and E) GIST-T1 cells were treated with 200 nM imatinib (IMA, KIT kinase inhibitor) for 8 hours. (D) Lysates were immunoblotted. (E) Proximity ligation assay (PLA) was performed with anti-KIT and anti-pPLCγ2^Y759^ antibodies. Distribution of Duolink PLA signals (green), Lectin-HPA (Golgi marker, red), and nucleus (DAPI, blue) are shown. Insets show magnified images of the Golgi region. Bars, 20 µm.

### PI4KIIIβ plays a key role in KIT retention as a downstream target of PKD2 in GIST cells

Next, we explored the downstream targets of PKD. Previous reports have shown that PI4P is generated when PKD activates PI4KIIIβ^22, 26, 37^, and then product PI4P recruits ADP-ribosylation factors (ARFs), ceramide transport protein (CERT), and oxysterol binding proteins (OSBPs) for proper membrane dynamics^22, 38, 39^. Importantly, PI4KIIIβ knockdown phenocopied PKD2 knockdown in both GIST-T1 (Figures 6A and 6B) and GIST430 (Figure S5B) cells. CERT knockdown showed similar results as PI4KIIIβ knockdown in KIT reduction and STAT5 dephosphorylation (Figure 6A). Given that the knockdown of PI4KIIIβ or CERT decreased KIT levels, the function of PKD2 in the control of membrane dynamics is important for mutant retention. In contrast, knockdown of OSBP1 or ARF1 did not affect KIT levels (Figure 6A). As OSBP and ARF consist of multiple members, a single knockdown may be unable to mimic the PKD2 knockdown phenotype. We next confirmed that PI4KIIIβ inhibitors (PI4KIIIβ-IN-10 and PIK-93) (ref. ^40^) decreased KIT, pAKT, and pSTAT5 levels like the PKD inhibitor (Figure 6C). Similar to the case of PKD2, PI4KIIIβ inhibition caused migration of KIT from the Golgi region to lysosomes (Figure 6D). As shown in Figure 6E, KIT was associated with PI4KIIIβ in a manner dependent on the mutant kinase activity. In support of these data, PI4KIIIβ localized together with KIT and PKD2 in the Golgi area (Figure 6F). Finally, we tested whether PI4KIIIβ lies downstream of KIT-PLCγ2-PKD2, by immunostaining for PI4P in IMA-or CRT-treated GIST-T1 cells. PI4P was detected in the Golgi area in untreated cells (Figure 6G, top panels), but the Golgi PI4P disappeared after treatment with IMA, CRT, or PIK-93 (Figure 6G), indicating that KIT retention in the Golgi area is dependent on PLCγ2-PKD2-PI4KIIIβ pathway activation.

**Figure 6.**
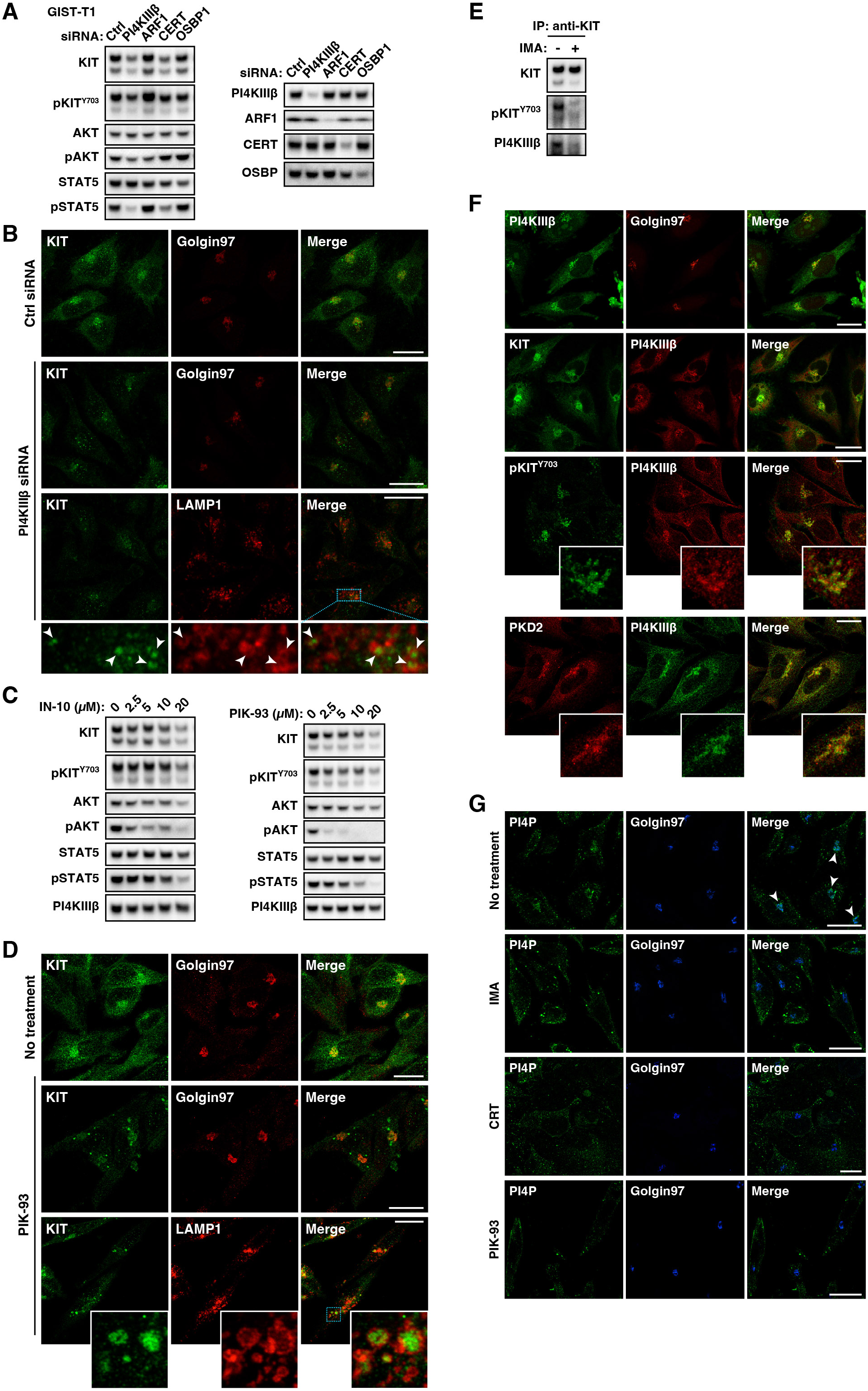
PI4KIIIβ plays a role in KIT retention downstream of PKD2. (A and B) GIST-T1 cells were transfected with indicated siRNAs for 30 hours. (A) Lysates were immunoblotted. (B) Cells were immunostained for KIT (green) in conjunction with golgin97 (red) or LAMP1 (lysosomal marker, red). Magnified images of boxed areas are shown. Bars, 20 µm. (C) GIST-T1 cells were treated with PI4KIIIβ inhibitors, IN-10 or PIK-93, for 16 hours and then immunoblotted. (D) GIST-T1 cells were treated with 20 µM PIK-93 for 4 hours and immunostained. Magnified images of boxed area are shown. Bars, 20 µm. (E) GIST-T1 cells were treated with 200 nM imatinib (IMA, KIT kinase inhibitor) for 8 hours. KIT was immunoprecipitated with an anti-KIT antibody and immunoprecipitates were immunoblotted. IP, immunoprecipitation. (F) GIST-T1 cells were immunostained using the indicated antibodies. Magnified images of the Golgi region. Bars, 20 µm. (G) GIST-T1 cells were treated with 200 nM IMA, 10 µM CRT0066101 (CRT, PKD inhibitor), or 20 µM PIK-93 for 16 hours. Cells were immunostained for PI4P (green) and golgin97 (blue). Arrowheads indicate the Golgi region. Bars, 20 µm. NB: IMA and CRT markedly decreased PI4P levels in the Golgi area.

### GGA1-AP1 complex plays a crucial role in the Golgi KIT retention in GIST cells

PI4P produced by PI4KIIIβ is believed to play many roles in the Golgi/TGN region, including recruitment of adaptors such as AP1 and GGA1-3 for the formation of clathrin-coated membrane carriers^26, 41, 42^. Therefore, we investigated whether these adaptor proteins were required for PKD2-induced KIT retention. Intriguingly, knockdown of γ-adaptin, an AP1 component, caused a substantial reduction in the KIT level, leading to inactivation of AKT and STAT5 in GIST-T1 cells (Figure 7A, left). Furthermore, knockdown of GGA1, but not GGA2/3, yielded similar results to γ-adaptin knockdown (Figure 7A, right). GGA1 knockdown decreased the protein levels of GGA2/3, suggesting that GGA1 is important for GGA2/3 stability. Next, we confirmed that KIT was taken in lysosomes in γ-adaptin-or GGA1-knocked down cells (Figures 7B and S6A). In GIST430 cells, γ-adaptin knockdown or GGA1 knockdown decreased KIT and pSTAT5 levels (Figure S6B). These adaptors were located in the Golgi area together with KIT, PKD2, and PI4KIIIβ (Figures 7C and S6C). To examine the extent of colocalization between these proteins in the Golgi/TGN region, we performed a quantitative analysis on signal overlapping between the PKD-related proteins and golgin97 (TGN)^43, 44^ or GOLPH4 (Golgi phosphoprotein 4, *medial-* Golgi) by calculating Pearson’s correlation coefficients. The super-resolution confocal live imaging microscopy (SCLIM) developed in RIKEN was used for this analysis^45–47^. Phospho-KIT, pPLCγ2, pPKD2, PI4KIIIβ, and γ-adaptin showed significantly higher correlation coefficients with golgin97 than GOLPH4 (Figures 7D and S6D), suggesting that KIT functionally associates with PKD2-related machineries in the *trans-*Golgi-to-TGN area.

**Figure 7.**
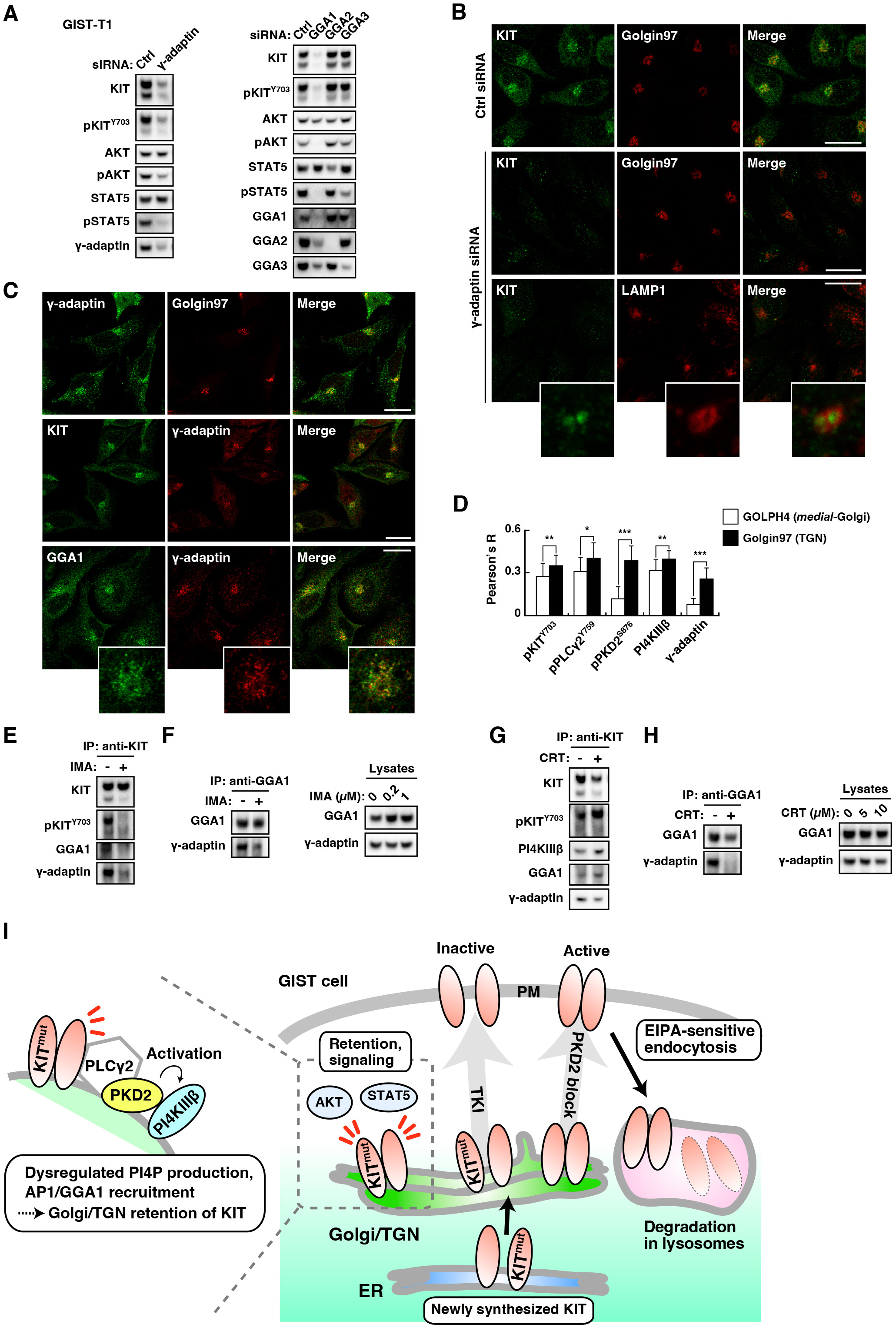
PI4KIIIβ effectors, γ-adaptin and GGA1, play a critical role in KIT retention in GIST cells. (A and B) GIST-T1 cells were transfected with the indicated siRNAs for 30 hours. (A) Lysates were immunoblotted. (B) Cells were immunostained for KIT (green) in conjunction with golgin97 (red) or LAMP1 (lysosomal marker, red). Magnified images of lysosomal region are shown. Bars, 20 µm. NB: Knockdown of γ-adaptin or GGA1 phenocopied that of PKD2 in KIT reduction and signal inhibition. (C and D) GIST-T1 cells were immunostained with the indicated antibodies. Magnified images of the perinuclear region are shown. Bars, 20 µm. (D) Pearson’s coefficients (Pearson’s R) were calculated by intensity correlation analysis of the indicated proteins vs. Golgi phosphoprotein 4 (GOLPH4, *medial*-Golgi protein) or golgin97 (*trans*-Golgi network (TGN) protein). Results are means ± s.d. (*n* = 11∼19). Asterisks indicate significant differences (\**P*<0.05; \*\**P*<0.01; \*\*\**P*<0.001) calculated by Student’s *t*-test. (E–H) GIST-T1 cells were treated with (E and F) 200 nM imatinib (IMA, KIT kinase inhibitor) or (G and H) 10 µM CRT0066101 (CRT, PKD inhibitor) for 8 hours. KIT (E and G) or GGA1 (F and H) was immunoprecipitated followed by immunoblotting. (I) In GIST cells, KIT is retained in the Golgi area during biosynthesis. KIT activates the PLCγ2-PKD2-PI4KIIIβ pathway to generate PI4P. Subsequently, PI4P recruits AP1 and GGA1 adaptors, and the complex retains KIT in the Golgi area (left side). Inhibition of this pathway by a PKD inhibitor, PKD2 knockdown, or tyrosine kinase inhibitors (TKIs), dissociates the KIT mutant from the Golgi region to the plasma membrane (PM). TKI such as imatinib blocks tyrosine kinase activity-dependent endocytosis, resulting in KIT retention on the PM. KIT activity is retained in the presence of PKD inhibitor, and the mutant undergoes ethyl-isopropyl amiloride-sensitive (EIPA-sensitive) endocytosis and is degraded in lysosomes. ER, endoplasmic reticulum.

Next, we performed a co-immunoprecipitation assay. Figure 7E shows that KIT was associated with γ-adaptin and GGA1 in a kinase-dependent manner. Furthermore, GGA1 was co-immunoprecipitated with γ-adaptin, and the interaction was decreased by IMA (Figure 7F). Considering our finding that both γ-adaptin knockdown and GGA1 knockdown caused migration of KIT from the Golgi region, the GGA1-AP1 interaction may be important for Golgi retention of KIT. Next, we investigated whether PKD inhibition affected the association between KIT and PKD2 effectors. Unexpectedly, PKD2, PI4KIIIβ, GGA1, and γ-adaptin were co-immunoprecipitated with KIT, even when the PKD2 activity was inhibited (Figure 7G). In contrast, PKD2 inhibition with CRT markedly decreased GGA1 association with γ-adaptin (Figure 7H). PKD activity is probably not essential for the association of KIT with PKD2, PI4KIIIβ, AP1, and GGA1, but is specifically required for the AP1-GGA1 interaction. Taken together, these results suggest that KIT recruits the AP1-GGA1 complex through the activation of PLCγ2-PKD2-PI4KIIIβ cascade for Golgi retention.

### Loss of function of PKD2-related machinery does not affect RTK levels and growth signaling in leukemia cells

Finally, we analyzed PKD in leukemia cells. M-07e (acute megakaryoblastic leukemia) expresses wild-type KIT, which is found on the PM, whereas the KIT mutant in HMC-1.1 (MCL), HMC-1.2, or Kasumi-1 cells (AML) localizes to endosome-lysosomal compartments^13–15, 48^ (see also Figure S7A). PKD1 expression in these leukemia cells was not detected by our immunoblotting. Moreover, PKD2 and PKD1/3 siRNA did not affect KIT levels or growth signaling (Figure S7B). Indeed, PKD2 was not co-immunoprecipitated with KIT in these cells (Figure S7C). Unlike in GIST cells, PLCγ2 knockdown in HMC-1.2 cells did not decrease KIT, pPKD2^S876^, or phospho-PKD substrates (Figure S7D). Furthermore, PKD2-related gene knockdown did not affect KIT levels or growth signals (Figures S7E and S7F). We were unable to detect an association between KIT and PI4KIIIβ, GGA1, or γ-adaptin (Figure S7G). GGA1 in HMC-1.2, as in GIST-T1, was in a complex with γ-adaptin, but the association was unaffected by PKD inhibition (Figure S7H), indicating that the regulation of GGA1-AP1 interaction in leukemia is different from that in GIST cells.

We recently reported that the FLT3-ITD accumulates in the Golgi area in AML cells, similarly to KIT in GIST cells^19^. However, PKD2 knockdown in the AML cell line, MOLM-14, did not affect FLT3 levels, signals, or mutant localization (Figures S8A and S8B). Therefore, PKD2 is not involved in the intracellular trafficking of RTK mutants in leukemia cells. Taken together, these results suggest that the PLCγ2-PKD2-PI4KIIIβ-AP1-GGA1 cascade plays a critical role in intracellular KIT retention specifically in GIST cells.

## DISCUSSION

In this study, we demonstrate that the KIT mutant is retained in the Golgi area in a manner dependent on PKD2 but not on PKD1/3 in GIST cells. In PKD2-inhibited GIST cells, KIT migrates from the Golgi region to the PM, and subsequently undergoes EIPA-sensitive endocytosis for lysosomal degradation (Figure 7I). We illustrate a model of a novel cascade in the Golgi area for KIT retention (Figure 7I, left): (i) KIT requires PLCγ2 for PKD2 activation, (ii) PI4KIIIβ is activated by PKD2 to generate PI4P, and (iii) the AP1-GGA1 complex is recruited by PI4P. These molecules are aberrantly recruited, presumably leading to KIT accumulation in the Golgi area. In sharp contrast, these PKD2-related proteins are not involved in the intracellular trafficking of KIT and FLT3-ITD in leukemia cells. Therefore, the PKD2-dependent retention of KIT mutants is a characteristic feature of GIST cells.

Although the requirement of PKD2-driven machinery for Golgi retention in KIT mutants is clear, how the mutant is retained in the Golgi area is still unclear. Recent studies showed that in yeast, vesicles budding from the TGN are recycled back to earlier Golgi cisternae in a manner dependent on AP1 (refs. ^49, 50^). In this context, AP1 might play a role in the intra-Golgi recycling of KIT in GIST cells. In contrast, the KIT mutant may be stably located in the Golgi/TGN membrane with the adaptor. Another possibility is that AP1-GGA1-dependent membrane carriers containing KIT rapidly bounce in and out of the TGN membrane. In addition, the physiological cargoes contained in PKD2-induced AP1 vesicles need to be investigated. Further studies are required to understand how KIT mutant hijacks physiological machinery, and to characterize PKD2-induced membrane carriers. New and more advanced tools including super-resolution live imaging in conjunction with fluorescent protein tagging technology and synchronization methods of early secretion, are required to understand local RTK dynamics in the Golgi/TGN, that will open new fields not only for oncology but also for cell biology.

AP1 and GGAs serve as clathrin adaptors for proper vesicular trafficking^51–53^. In addition to these studies, our data show that AP1 associates with GGA1 in both GIST and MCL. PKD2-induced AP1-GGA1 interaction, however, traps KIT in the Golgi specifically in GIST cells. Whether AP1-GGA1 vesicles are involved in physiological intracellular trafficking remains unknown, however our study suggests a possibility that some vesicles may contain multiple adaptor subsets.

Unexpectedly, FLT3-ITD is retained in the Golgi area in AML cells in a PKD2-independent manner, and knockdown of PKD1, PKD2, or PKD3 does not affect FLT3 signals, indicating that the cause of Golgi retention in FLT3-ITD is not PKD activation. Golgi retention of individual RTKs is regulated by different molecular mechanisms. Previous studies showed that cancer-causing RTKs, such as activated PDGFRA, FGFR3, TRKA, IGF-1, and MET, are also found in the Golgi area^54–61^. Additionally, non-receptor type signaling molecules, such as SRC-family kinases and RAS, accumulate in the Golgi area^16, 62–65^. These Golgi retentions may or may not depend on PKD2 activity. Thus, further studies are required to understand the mechanism of Golgi retention of individual oncogenic signaling molecules.

Recent studies have shown that loss-of-function mutations of coatomer subunit α (COPA), which plays a role in intra-Golgi transport as a member of the coat protein complex I (COPI), cause retention and excessive activation of STING (stimulator of IFN genes) in the Golgi area, resulting in autoinflammation called COPA syndrome^66–68^. Mutations in AP1 subunits, such as σ-adaptins, lead to the development of mental retardation, presumably through dysregulated vesicular trafficking^69, 70^. Thus, elucidating the homeostatic mechanisms of adaptor proteins in the Golgi/TGN is important to understand the etiology of these diseases. In this study, we observed that mutant KIT affects adaptor protein function suggesting that aberrantly activated RTK in the Golgi/TGN may dysregulate membrane traffic machinery. These findings raise the intriguing possibility that RTKs activated via gene amplification or mutation affect the distribution of membrane proteins related to cancer development. Because specific PM proteins are involved in immune escape and invasion, understanding the role of cancer-causing RTKs in the Golgi/TGN will help improve cancer treatments.

Molecular-targeting drugs, such as TKIs, for solid tumors with RTK mutations significantly extend the lifespan of patients^71^. However, most tumors acquire TKI resistance within a couple of years^10, 71^. In most GISTs, *KIT ex13* or *ex17* mutations are frequently found as TKI-resistant alterations^10–12^. In this study, pharmacological blockade of PKD2 or PI4KIIIβ suppressed KIT signaling by normalizing receptor trafficking from the Golgi region, regardless of the site of *KIT* mutation. PKD and PI4KIIIβ inhibitors are used in preclinical trials for the treatment of solid tumors^72–77^, thus investigating these compounds in animal models for GISTs will drive progress in the development of enhanced cancer treatments. Previously, we developed novel strategies for suppressing oncogenic Golgi-localized RTK signaling including (i) blocking RTK trafficking from the ER to the Golgi^14, 15, 17, 19^, and (ii) disruption of RTK stability in organelles through HSP90 inhibition^18^. Our previous findings and this study suggest that understanding the mechanism of intracellular trafficking and the induction of degradation will help us establish a novel method for the suppression of cancer cell proliferation. Additionally, this study provides a new strategy for the inhibition of TKI-resistant KIT mutants in GISTs through dissociation of the mutant from the signaling platform and subsequent induction of lysosomal KIT degradation.

The three PKD members are structurally similar^30, 31^, and are expressed in GIST cells, but only PKD2 functions in the Golgi retention of KIT. PKD2, but not PKD1 and 3, has an amino-terminal proline-rich and a serine-rich domain^30, 32, 38^, which could confer specific functions to PKD2. Structural and functional analyses of PKD2 mutants lacking the proline-and serine-rich domains or insertion of these domains into PKD1 and 3 will help understand PKD2-specific functions in KIT retention. Our future studies will further examine the structure-function relationship of PKD2 in GIST. Furthermore, we have identified PKD2 as a downstream target of KIT in the Golgi/TGN. We hypothesize that PKD2 activated by KIT in the Golgi area can phosphorylate signaling molecules, even if they are not involved in intracellular trafficking. Previous reports have shown that PKD2 directly phosphorylates cortactin and CIB1 (calcium and integrin binding 1), resulting in cell migration and angiogenesis, respectively^33, 78, 79^. It would be of great interest to examine whether KIT activates these proteins via PKD2 activation in the Golgi/TGN in GIST cells.

In conclusion, we show that KIT mutants are selectively retained in the Golgi area through aberrant involvement of the PLCγ2-PKD2-PI4KIIIβ-AP1-GGA1 cascade in GIST cells. These observations provide new insights into the significance of mislocalization of RTK mutants as well as of other mutant signaling molecules. Moreover, from a clinical point of view, our findings offer a new strategy for the inhibition of oncogenic signaling by releasing RTK from the Golgi/TGN signaling platform toward lysosomes. Therefore, our study provides a significant insight into intracellular trafficking in molecular cell biology and also for oncology and targeted therapeutics.

## MATERIALS & METHODS

### Cell culture

GIST-T1 (Cosmo Bio, Tokyo, Japan) and GIST-R9 cells were cultured at 37°C in DMEM supplemented with 10% fetal calf serum (FCS), penicillin, and streptomycin (Pen/Strep). GIST-R9 cells were maintained in the presence of 1 µM IMA as previously described^17, 25^. GIST430/654 (hereafter referred to as GIST430) and GIST48 were kindly provided by Dr. Jonathan Fletcher (Dana-Farber Cancer Institute, Boston, MA)^24^. GIST430 and GIST48 cells were cultured at 37°C in IMDM supplemented with 15% FCS and Pen/Strep. GIST430 cells were maintained in the presence of 100 nM IMA. Kasumi-1 (JCRB Cell Bank, Osaka, Japan), M-07e, MOLM-14 (DSMZ, Braunschweig, Germany), HMC-1.2, and HMC-1.1 cells (Merck, Darmstadt, Germany) were cultured at 37°C in RPMI1640 supplemented with 10% FCS, Pen/Strep, and 50 µM 2-mercaptoethanol. For expansion of M-07e, 10 ng/mL GM-CSF (Peprotech, Rocky Hill, NJ, USA) was used. All human cell lines were authenticated by STR analysis at the JCRB Cell Bank and tested for *Mycoplasma* contamination using a MycoAlert Mycoplasma Detection Kit (Lonza, Basel, Switzerland).

### Antibodies and Chemicals

The list of antibodies used for immunoblotting and immunofluorescence is shown in Supplementary Tables 1–3. A list of small compounds with their sources is provided in Supplementary Table 4.

### Gene silencing with siRNA

To silence *PKD*-related genes, ON-TARGETplus SMARTpool siRNAs were purchased from Dharmacon (Lafayette, CO, USA). A list of the siRNAs used is provided in Supplementary Table 5. Electroporation was performed using the NEON Transfection System (Thermo Fisher Scientific, Rockford, IL, USA), according to the manufacturer’s instructions.

### Immunofluorescence confocal microscopy

GIST cells were cultured on poly L-lysine-coated coverslips and fixed with 4% paraformaldehyde (PFA) for 20 min at room temperature. Leukemia cells in suspension culture were fixed with MeOH at-20°C for at least 10 min or with 4% PFA for 20 min at room temperature and then cyto-centrifuged onto coverslips. Fixed cells were permeabilized and blocked for 30 min in D-PBS(-) supplemented with 0.1% saponin and 3% BSA and then incubated with primary and secondary antibodies for 1 h each. After washing with D-PBS(-), the cells were mounted with Fluoromount (DiagnosticBioSystems, Pleasanton, CA, USA). Confocal images were obtained with a Fluoview FV10i (Olympus, Tokyo, Japan) or TCS SP5 II/SP8 (Leica, Wetzlar, Germany) laser scanning microscope. For super-resolution imaging, a custom-made confocal microscopic system (super-resolution confocal live imaging microscopy, SCLIM) developed in RIKEN was used^45–47^. Composite figures were prepared using FV1000 Viewer (Olympus), LAS X (Leica), Volocity (PerkinElmer, Waltham, MA), Photoshop, and Illustrator software (Adobe, San Jose, CA, USA).

### Proximity ligation assay (PLA)

For PLA, the Duolink^TM^ *In Situ* Green Starter Kit Mouse/Rabbit (Sigma-Aldrich, St. Louis, MO, USA) was used. GIST-T1 cells were fixed, permeabilized, and stained with primary antibodies as described above. Subsequently, PLA probes, anti-mouse PLUS, and anti-rabbit MINUS were incubated with the cells for 1 h at 37°C. After washing, the PLUS and MINUS strand DNAs were circulated and amplified to form a chromophore according to the manufacturer’s instructions. To visualize the Golgi area without an antibody, lectin-HPA was used. Nuclei were stained with DAPI in mounting medium.

### Immunostaining for PI4P visualization

GIST-T1 cells were treated with 50 mM NH_4_Cl in D-PBS(−) before fixation with 2% PFA^80^. Permeabilization and staining were performed as described above. To stain for PI4P, anti-PI4P mouse IgM and Alexa Fluor-488-anti-mouse IgM were used as the primary and secondary antibodies, respectively. After washing with D-PBS(-), stained cells were fixed with 2% PFA for 15 min, treated with 50 mM NH_4_Cl in D-PBS(-), and washed with water.

### Western blotting

Lysates prepared in SDS-PAGE sample buffer were subjected to SDS-PAGE and electrotransferred onto polyvinylidene fluoride membranes. Briefly, 5% skim milk in TBS-T was used to dilute the antibodies. For immunoblotting with anti-KIT [pY703], anti-PLCγ2 [pY759], or anti-pTyr, the antibodies were diluted with 3% BSA in TBS-T. Immunodetection was performed using Immobilon Western Chemiluminescent HRP Substrate (Sigma-Aldrich). Sequential re-probing of membranes was performed after complete removal of antibodies in stripping buffer (Thermo Fisher Scientific) or inactivation of peroxidase by 0.1% NaN_3_. The results were analyzed using ChemiDoc XRC+ with Image Lab software (Bio-Rad, Hercules, CA, USA).

### Immunoprecipitation

Lysates from 2–4 x 10^6^ cells were prepared in NP-40 lysis buffer (50 mM HEPES pH 7.4, 10% glycerol, 1% NP-40, 4 mM EDTA, 100 mM NaF, 1 mM Na_3_VO_4_, protease inhibitor cocktail, 2 mM β-glycerophosphate, 2 mM sodium pyrophosphate, and 1 mM PMSF). Immunoprecipitation was performed at 4°C for 5–8 hours using protein G Sepharose or Dynabeads precoated with antibodies. The immunoprecipitates were dissolved in SDS-PAGE sample buffer.

### Statistical analyses

Differences between two groups were analyzed by the two-tailed Student’s *t*-test. *P* values are described in the legend for Figure 7 (\**P* < 0.05, \*\**P* < 0.01, \*\*\**P* < 0.001).

## ABBREVIATIONS

AML: acute myelogenous leukemia
AP1: adaptor protein 1
ARF: ADP-ribosylation factor
CERT: ceramide transport protein
CQ: chloroquine
CRT: CRT0066101
EIPA: ethyl-isopropyl amiloride
ER: endoplasmic reticulum
ex: exon
FLT3-ITD: FMS-like tyrosine kinase 3-intenal tandem duplication
GGA: Golgi-associated, γ-adaptin ear-containing, ARF-binding protein
GIST: gastrointestinal stromal tumor
mut: mutant
GM130: Golgi matrix protein 130 kDa
GOLPH4: Golgi phosphoprotein 4
IMA: imatinib
IP/IPs: immunoprecipitation/immunoprecipitates
LAMP1: lysosomal-associated membrane protein 1
MCL: mast cell leukemia
NH_4_Cl: ammonium chloride
OSBP: oxysterol binding proteins
PDGFR: platelet-derived growth factor receptor
PI4K: phosphatidylinositol 4-kinase
PI4P: phosphatidylinositol 4-phosphate
PKD: protein kinase D
pKIT: phospho-KIT
PLA: proximity ligation assay
PLC: phospholipase C
PM: plasma membrane
RTK: receptor tyrosine kinase
SCLIM: super-resolution confocal live imaging microscopy
siRNA: small interfering RNA
STAT: signal transducer and activator of transcription
TGN: *trans-*Golgi network
TKI: tyrosine kinase inhibitor
wt: wild-type

## ACKNOWLEDGMENTS

The authors thank Ms. Kumiko Ishii (RIKEN) for maintaining GIST-T1 cells. This work was supported by a grant-in-aid for Scientific Research from the Japan Society for the Promotion of Science (21K07163 to YO, 20K08719 to RA, and 22K08883 to TN), a research grant from the Takeda Science Foundation (to YO), the Research Foundation for Pharmaceutical Sciences (to YO), and the Okinaka Memorial Institute for Medical Research (to YO).

## AUTHOR CONTRIBUTIONS

Y.O. conceived, designed, performed, and analyzed the data from all experiments and wrote the manuscript. K.K. performed confocal laser scanning microscopic analysis. T. Tojima helped analyze the microscopy data. M.N. performed the immunoblotting. I.S. analyzed the data and edited the manuscript. T. Takahashi provided advice on the design of the *in vitro* experiments for IMA-resistant GIST cells. R.A. provided advice on the design of the *in vitro* experiments for AML cells. A.N. analyzed the data and edited the manuscript. T.N. conceived and supervised the project, analyzed the data, and wrote the manuscript. All authors have read and approved the final manuscript.

## REFERENCES

1. Lemmon, M.A. & Schlessinger, J. Cell signaling by receptor tyrosine kinases. Cell 141, 1117–1134 (2010). https://doi.org/10.1016/j.cell.2010.06.011.

2. Thomsen, L. et al. Interstitial cells of Cajal generate a rhythmic pacemaker current. Nat. Med. 4, 848–851 (1998). https://doi.org/10.1038/nm0798-848.

3. Lennartsson, R. & Rönnstrand, L. Stem cell factor receptor/c-Kit: from basic science to clinical implications. Physiol. Rev. 92, 1619–1649 (2012). https://doi.org/10.1152/physrev.00046.2011.

4. Roskoski, R. Structure and regulation of Kit protein-tyrosine kinase - the stem cell factor receptor. Biochem. Biophys. Res. Commun. 338, 1307–1315 (2005). https://doi.org/10.1016/j.bbrc.2005.09.150.

5. Thömmes, K., Lennartsson, J., Carlberg, M. & Rönnstrand, L. Identification of Tyr-703 and Tyr-936 as the primary association sites for Grb2 and Grb7 in the c-Kit/stem cell factor receptor. Biochem. J. 341, 211–216 (1999). https://doi.org/10.1042/bj3410211.

6. Hirota, S. et al. Gain-of-function mutations of *c-kit* in human gastrointestinal stromal tumors. Science 279, 577–580 (1998). https://doi.org/10.1126/science.279.5350.577.

7. Nishida, T. et al. Familial gastrointestinal stromal tumours with germline mutation of the *KIT* gene. Nat. Genet. 19, 323–324 (1998). https://doi.org/10.1038/1209.

8. Kitamura, Y. Gastrointestinal stromal tumors: past, present, and future. J. Gastroenterol. 43, 499–508 (2008). https://doi.org/10.1007/s00535-008-2200-y.

9. Boissan, M., Feger, F., Guillosson, J.J. & Arock, M. c-Kit and c-kit mutations in mastocytosis and other hematological diseases. J. Leukoc. Biol. 67, 135–148 (2000). https://doi.org/10.1002/jlb.67.2.135.

10. Blay, J.Y., Kang, Y.K., Nishida, T. & von Mehren, M. Gastrointestinal stromal tumours. Nat. Rev. Dis. Primers 7, 22 (2021). https://doi.org/10.1038/s41572-021-00254-5.

11. Lasota, J. & Miettinen, M. Clinical significance of oncogenic *KIT* and *PDGFRA* mutations in gastrointestinal stromal tumours. Histopathology 53, 245–266 (2008). https://doi.org/10.1111/j.1365-2559.2008.02977.x.

12. Nishida, T. et al. Secondary mutations in the kinase domain of the *KIT* gene are predominant in imatinib-resistant gastrointestinal stromal tumor. Cancer Sci. 99, 799–804 (2008). https://doi.org/10.1111/j.1349-7006.2008.00727.x.

13. Obata, Y. et al. Oncogenic Kit signals on endolysosomes and endoplasmic reticulum are essential for neoplastic mast cell proliferation. Nat. Commun. 5, 5715 (2014). https://doi.org/10.1038/ncomms6715.

14. Hara, Y. et al. M-COPA suppresses endolysosomal Kit-Akt oncogenic signalling through inhibiting the secretory pathway in neoplastic mast cells. PLoS ONE 12, e0175514 (2017). https://doi.org/10.1371/journal.pone.0175514.

15. Obata, Y. et al. N822K-or V560G-mutated KIT activation occurs preferentially in lipid rafts of the Golgi apparatus in leukemia cells. Cell Commun. Signal. 17, 114 (2019). https://doi.org/10.1186/s12964-019-0426-3.

16. Obata, Y. et al. Oncogenic signaling by Kit tyrosine kinase occurs selectively on the Golgi apparatus in gastrointestinal stromal tumors. Oncogene 36, 3661–3672 (2017). https://doi.org/10.1038/onc.2016.519.

17. Obata, Y. et al. Oncogenic Kit signalling on the Golgi is suppressed by blocking secretory trafficking with M-COPA in GISTs. Cancer Lett. 415, 1–10 (2018). https://doi.org/10.1016/j.canlet.2017.11.032.

18. Saito, Y. et al. TAS-116 inhibits oncogenic KIT signalling on the Golgi in both imatinib-naïve and imatinib-resistant gastrointestinal stromal tumours. Br. J. Cancer 122, 658–667 (2020). https://doi.org/10.1038/s41416-019-0688-y.

19. Yamawaki, K. et al. FLT3-ITD transduces autonomous growth signals during its biosynthetic trafficking in acute myelogenous leukemia cells. Sci. Rep. 11, 22678 (2021). https://doi.org/10.1038/s41598-021-02221-2.

20. Liljedahl, M. et al. Protein kinase D regulates the fission of cell surface destined transport carriers from the trans-Golgi network. Cell 104, 409–420 (2001). https://doi.org/10.1016/S0092-8674(01)00228-8.

21. Baron, C.L. & Malhotra, V. Role of diacylglycerol in PKD recruitment to the TGN and protein transport to the plasma membrane. Science 295, 325–328 (2002). https://doi.org/10.1126/science.1066759.

22. Malhotra, V. & Campelo, F. PKD regulates membrane fission to generate TGN to cell surface transport carriers. Cold Spring Harb. Perspect. Biol. 3, a005280 (2011). https://doi.org/10.1101/cshperspect.a005280.

23. Taguchi, T. et al. Conventional and molecular cytogenetic characterization of a new human cell line, GIST-T1, established from gastrointestinal stromal tumor. Lab. Invest. 82, 663–665 (2002). https://doi.org/10.1038/labinvest.3780461.

24. Bauer, S., Duensing, A., Demetri, G.D. & Fletcher, J.A. KIT oncogenic signaling mechanisms in imatinib-resistant gastrointestinal stromal tumor: PI3-kinase/AKT is a crucial survival pathway. Oncogene 26, 7560–7568 (2007). https://doi.org/10.1038/sj.onc.1210558.

25. Takahashi, T. et al. Genomic and transcriptomic analysis of imatinib resistance in gastrointestinal stromal tumors. Genes Chromosomes Cancer 56, 303–313 (2017). https://doi.org/10.1002/gcc.22438.

26. Bankaitis, V.A., Garcia-Mata, T. & Mousley, C.J. Golgi membrane dynamics and lipid metabolism. Curr. Biol. 22, R414–424 (2012). https://doi.org/10.1016/j.cub.2012.03.004.

27. Wille, C. et al. Protein kinase D2 induces invasion of pancreatic cancer cells by regulating matrix metalloproteinases. Mol. Biol. Cell. 25, 324–336 (2014). https://doi.org/10.1091/mbc.e13-06-0334.

28. Gekle, M. et al. Inhibition of Na^+^-H^+^ exchange impairs receptor-mediated albumin endocytosis in renal proximal tubule-derived epithelial cells from opossum. J. Physiol. 520, 709–721 (1999). https://doi.org/10.1111/j.1469-7793.1999.00709.x.

29. Recouvreux, M.V. & Commisso, C. Macropinocytosis: A Metabolic Adaptation to Nutrient Stress in Cancer. Front. Endocrinol. 8, 261 (2017). https://doi.org/10.3389/fendo.2017.00261.

30. Van Lint, J. et al. Protein kinase D: an intracellular traffic regulator on the move. Trends Cell Biol. 12, 193–200 (2002). https://doi.org/10.1016/S0962-8924(02)02262-6.

31. Roy, A., Ye, J., Deng, F. & Wang, Q.J. Protein kinase D signaling in cancer: A friend or foe? Biochim. Biophys. Acta Rev. Cancer 1868, 283–294 (2017). https://doi.org/10.1016/j.bbcan.2017.05.008.

32. Azoitei, N., Cobbaut, M., Becher, A., Van Lint, J. & Seufferlein, T. Protein kinase D2: a versatile player in cancer biology. Oncogene 37, 1263–1278 (2018). https://doi.org/10.1038/s41388-017-0052-8.

33. Weeber, F., Becher, A., Seibold, T., Seufferlein, T. & Eiseler, T. Concerted regulation of actin polymerization during constitutive secretion by cortactin and PKD2. J. Cell Sci. 132, jcs232355 (2019). https://doi.org/10.1242/jcs.232355.

34. Cooke, M. & Kazanietz, M.G. Overarching roles of diacylglycerol signaling in cancer development and antitumor immunity. Sci. Signal. 15, eabo0264d (2022). https://doi.org/10.1126/scisignal.abo0264.

35. Carpenter, G. & Ji, Q. Phospholipase C-gamma as a signal-transducing element. Exp. Cell Res. 253, 15–24 (1999). https://doi.org/10.1006/excr.1999.4671.

36. Obeng, E.O. et al. Phosphoinositide-Dependent Signaling in Cancer: A Focus on Phospholipase C Isozymes. Int. J. Mol. Sci. 21, 2581 (2020). https://doi.org/10.3390/ijms21072581.

37. Hausser, A. et al. Protein kinase D regulates vesicular transport by phosphorylating and activating phosphatidylinositol-4 kinase IIIβ at the Golgi complex. Nat. Cell Biol. 7, 880–886 (2005). https://doi.org/10.1038/ncb1289.

38. Pusapati, G.V. et al. Role of the Second Cysteine-rich Domain and Pro275 in Protein Kinase D2 Interaction with ADP-Ribosylation Factor 1, *Trans*-Golgi Network Recruitment, and Protein Transport. Mol. Biol. Cell 21, 1011–1022 (2010). https://doi.org/10.1091/mbc.e09-09-0814.

39. Roussel, É. & Lippé, L. Cellular Protein Kinase D Modulators Play a Role during Multiple Steps of Herpes Simplex Virus 1 Egress. J. Virol. 92, e01486–18 (2018). https://doi.org/10.1128/JVI.01486-18.

40. Boura, E. & Nencka, R. Phosphatidylinositol 4-kinases: Function, structure, and inhibition. Exp. Cell Res. 337, 136–145 (2015). https://doi.org/10.1016/j.yexcr.2015.03.028.

41. Wang, Y.J. et al. Phosphatidylinositol 4 phosphate regulates targeting of clathrin adaptor AP-1 complexes to the Golgi. Cell 114, 299–310 (2003). https://doi.org/10.1016/S0092-8674(03)00603-2.

42. Wang, J. et al. PI4P Promotes the Recruitment of the GGA Adaptor Proteins to the *Trans*-Golgi Network and Regulates Their Recognition of the Ubiquitin Sorting Signal. Mol. Biol. Cell 18, 2646–2655 (2007). https://doi.org/10.1091/mbc.e06-10-0897.

43. Tie, H.C., Ludwig, A., Sandin, S. & Lu, L. The spatial separation of processing and transport functions to the interior and periphery of the Golgi stack. eLife 7, e41301 (2018). https://doi.org/10.7554/eLife.41301.

44. Tie, H.C., Mahajan, D. & Lu, L. Visualizing intra-Golgi localization and transport by side-averaging Golgi ministacks. J. Cell Biol. 221, e202109114 (2022). https://doi.org/10.1083/jcb.202109114.

45. Kurokawa, K., Ishii, M., Suda, Y., Ichihara, A. & Nakano, A. Live cell visualization of Golgi membrane dynamics by super-resolution confocal live imaging microscopy. Methods Cell Biol. 118, 235–242 (2013). https://doi.org/10.1016/B978-0-12-417164-0.00014-8.

46. Kurokawa, K. et al. Visualization of secretory cargo transport within the Golgi apparatus. J Cell Biol. 218, 1602–1618 (2019). https://doi.org/10.1083/jcb.201807194.

47. Kurokawa, K. & Nakano, A. Live-cell imaging by super-resolution confocal live imaging microscopy (SCLIM): simultaneous three-color and four-dimensional live cell imaging with high space and time resolution. Bio-Protocol 10, e3732 (2020). https://doi.org/10.21769/BioProtoc.3732.

48. Rai, S. et al. Chlorpromazine eliminates acute myeloid leukemia cells by perturbing subcellular localization of FLT3-ITD and KIT-D816V. Nat. Commun. 11, 4147 (2020). https://doi.org/10.1038/s41467-020-17666-8.

49. Casler, J.C., Papanikou, E., Barrero, J.J. & Glick, B.S. Maturation-driven transport and AP-1-dependent recycling of a secretory cargo in the Golgi. J. Cell Biol. 218, 1582–1601 (2019). https://doi.org/10.1083/jcb.201807195.

50. Casler, J.C. et al. Clathrin adaptors mediate two sequential pathways of intra-Golgi recycling. J. Cell Biol. 221, e202103199 (2022). https://doi.org/10.1083/jcb.202103199.

51. Doray, B., Ghosh, P., Griffith, J., Geuze, H.J. & Kornfeld, S. Cooperation of GGAs and AP-1 in packaging MPRs at the trans-Golgi network. Science 297, 1700–1703 (2002). https://doi.org/10.1126/science.1075327.

52. Bai, H., Doray, B. & Kornfeld, S. GGA1 interacts with the adaptor protein AP-1 through a WNSF sequence in its hinge region. J. Biol. Chem. 279, 17411–17417 (2004). https://doi.org/10.1074/jbc.M401158200.

53. Hirst, J. et al. Distinct and overlapping roles for AP-1 and GGAs revealed by the “knocksideways” system. Curr. Biol. 22, 1711–1716 (2012). https://doi.org/10.1016/j.cub.2012.07.012.

54. Ronchetti, D. et al. Deregulated FGFR3 mutants in multiple myeloma cell lines with t(4;14): comparative analysis of Y373C, K650E and the novel G384D mutations. Oncogene 20, 3553–3562 (2001). https://doi.org/10.1038/sj.onc.1204465.

55. Gibbs, L. & Legeai-Maller, L. FGFR3 intracellular mutations induce tyrosine phosphorylation in the Golgi and defective glycosylation. Biochim. Biophys. Acta. 1773, 502–512 (2007). https://doi.org/10.1016/j.bbamcr.2006.12.010.

56. Bahlawane, C. et al. Constitutive activation of oncogenic PDGFRα-mutant proteins occurring in GIST patients induces receptor mislocalisation and alters PDGFRα signalling characteristics. Cell Commun. Signal. 13, 21 (2015). https://doi.org/10.1186/s12964-015-0096-8.

57. Farina, A.R. et al. Retrograde TrkAIII transport from ERGIC to ER: a re-localisation mechanism for oncogenic activity. Oncotarget 6, 35636–35651 (2015). https://doi.org/10.18632/oncotarget.5802.

58. Ip, C.K.M. et al. Neomorphic PDGFRA extracellular domain driver mutations are resistant to PDGFRA targeted therapies. Nat. Commun. 9, 4583 (2018). https://doi.org/10.1038/s41467-018-06949-w.

59. Frazier, N.M., Brand, T., Gordan, J.D., Grandis, J. & Jura, N. Overexpression-mediated activation of MET in the Golgi promotes HER3/ERBB3 phosphorylation. Oncogene 38, 1936–1950 (2018). https://doi.org/10.1038/s41388-018-0537-0.

60. Schmidt-Arras, D. & Böhmer, F.D. Mislocalisation of Activated Receptor Tyrosine Kinases-Challenges for Cancer Therapy. Trends Mol. Med. 26, 833–847 (2020). https://doi.org/10.1016/j.molmed.2020.06.002.

61. Rieger, L. et al. IGF-1 receptor activity in the Golgi of migratory cancer cells depends on adhesion-dependent phosphorylation of Tyr^1250^ and Tyr^1251^. Sci. Signal. 13, eaba3176 (2020). https://doi.org/10.1126/scisignal.aba3176.

62. Chiu, V.K. et al. Ras signalling on the endoplasmic reticulum and the Golgi. Nat. Cell Biol. 4, 343–350 (2001). https://doi.org/10.1038/ncb783.

63. Rocks, O. et al. An acylation cycle regulates localization and activity of palmitoylated Ras isoforms. Science 307, 1746–1752 (2005). https://doi.org/10.1126/science.1105654.

64. Pulvirenti, T. et al. A traffic-activated Golgi-based signalling circuit coordinates the secretory pathway. Nat. Cell Biol. 10, 912–922 (2008). https://doi.org/10.1038/ncb1751.

65. Sato, I. et al. Differential trafficking of Src, Lyn, Yes and Fyn is specified by the state of palmitoylation in the SH4 domain. J. Cell Sci. 122, 965–975 (2009). https://doi.org/10.1242/jcs.034843.

66. Watkin, L.B. et al. *COPA* mutations impair ER-Golgi transport and cause hereditary autoimmune-mediated lung disease and arthritis. Nat. Genet. 47, 654–660 (2015). https://doi.org/10.1038/ng.3279.

67. Lepelley, A. et al. Mutations in *COPA* lead to abnormal trafficking of STING to the Golgi and interferon signaling. J. Exp. Med. 217, e20200600 (2020). https://doi.org/10.1084/jem.20200600.

68. Steiner, A. et al. Deficiency in coatomer complex I causes aberrant activation of STING signalling. Nat. Commun. 13, 2321 (2022). https://doi.org/10.1038/s41467-022-29946-6.

69. Tarpey, P.S. et al. Mutations in the gene encoding the Sigma 2 subunit of the adaptor protein 1 complex, AP1S2, cause X-linked mental retardation. Am. J. Hum. Genet. 79, 1119–1124 (2006). https://doi.org/10.1086/510137.

70. Montpetit, A., et al. Disruption of AP1S1, causing a novel neurocutaneous syndrome, perturbs development of the skin and spinal cord. PLoS Genet. 4, e1000296 (2008). https://doi.org/10.1371/journal.pgen.1000296.

71. Schmitt, M.W., Loeb, L.A. & Salk, J.J. The influence of subclonal resistance mutations on targeted cancer therapy. Nat. Rev. Clin. Oncol. 13, 335–347 (2016). https://doi.org/10.1038/nrclinonc.2015.175.

72. Harikumar, K.B. et al. Anovel small-molecule inhibitor of protein kinase D blocks pancreatic cancer growth *in vitro* and *in vivo*. Mol. Cancer Ther. 9, 1136–1146 (2010). https://doi.org/10.1158/1535-7163.MCT-09-1145.

73. Wei, N., Chu, E., Wipf, P. & Schmitz, J.C. Protein kinase D as a potential chemotherapeutic target for colorectal cancer. Mol. Cancer Ther. 13, 1130–1141 (2014). https://doi.org/10.1158/1535-7163.MCT-13-0880.

74. Borges, S. et al. Effective Targeting of Estrogen Receptor-Negative Breast Cancers with the Protein Kinase D Inhibitor CRT0066101. Mol. Cancer Ther. 14, 1306–1316 (2015). https://doi.org/10.1158/1535-7163.MCT-14-0945.

75. Tan, X. et al. PI4KIIIβ is a therapeutic target in chromosome 1q-amplified lung adenocarcinoma. Sci. Transl. Med. 12, eaax3772 (2020). https://doi.org/10.1126/scitranslmed.aax3772.

76. Lv, D. et al. Small-Molecule Inhibitor Targeting Protein Kinase D: A Potential Therapeutic Strategy. Front. Oncol. 11, 680221 (2021). https://doi.org/10.3389/fonc.2021.680221.

77. Shi, L. et al. Addiction to Golgi-resident PI4P synthesis in chromosome 1q21.3-amplified lung adenocarcinoma cells. Proc. Natl Acad. Sci. USA 118, e2023537118 (2021). https://doi.org/10.1073/pnas.2023537118.

78. Eiseler, T., Hausser, A., De Kimpe, L., Van Lint, J. & Pfizenmaier, K. Protein kinase D controls actin polymerization and cell motility through phosphorylation of cortactin. J. Biol. Chem. 285, 18672–18683 (2010). https://doi.org/10.1074/jbc.M109.093880.

79. Armacki, M. et al. A novel splice variant of calcium and integrin-binding protein 1 mediates protein kinase D2-stimulated tumour growth by regulating angiogenesis. Oncogene 33, 1167–1180 (2014). https://doi.org/10.1038/onc.2013.43.

80. Capasso, S. et al. Sphingolipid metabolic flow controls phosphoinositide turnover at the *trans-*Golgi network. EMBO J. 36, 1736–1754 (2017). https://doi.org/10.15252/embj.201696048.

