## Supplementary Figures for "Golgi retention and oncogenic KIT signaling via PLCγ2-PKD2-PI4KIIIβ activation in GIST cells"

### Supplemental figures

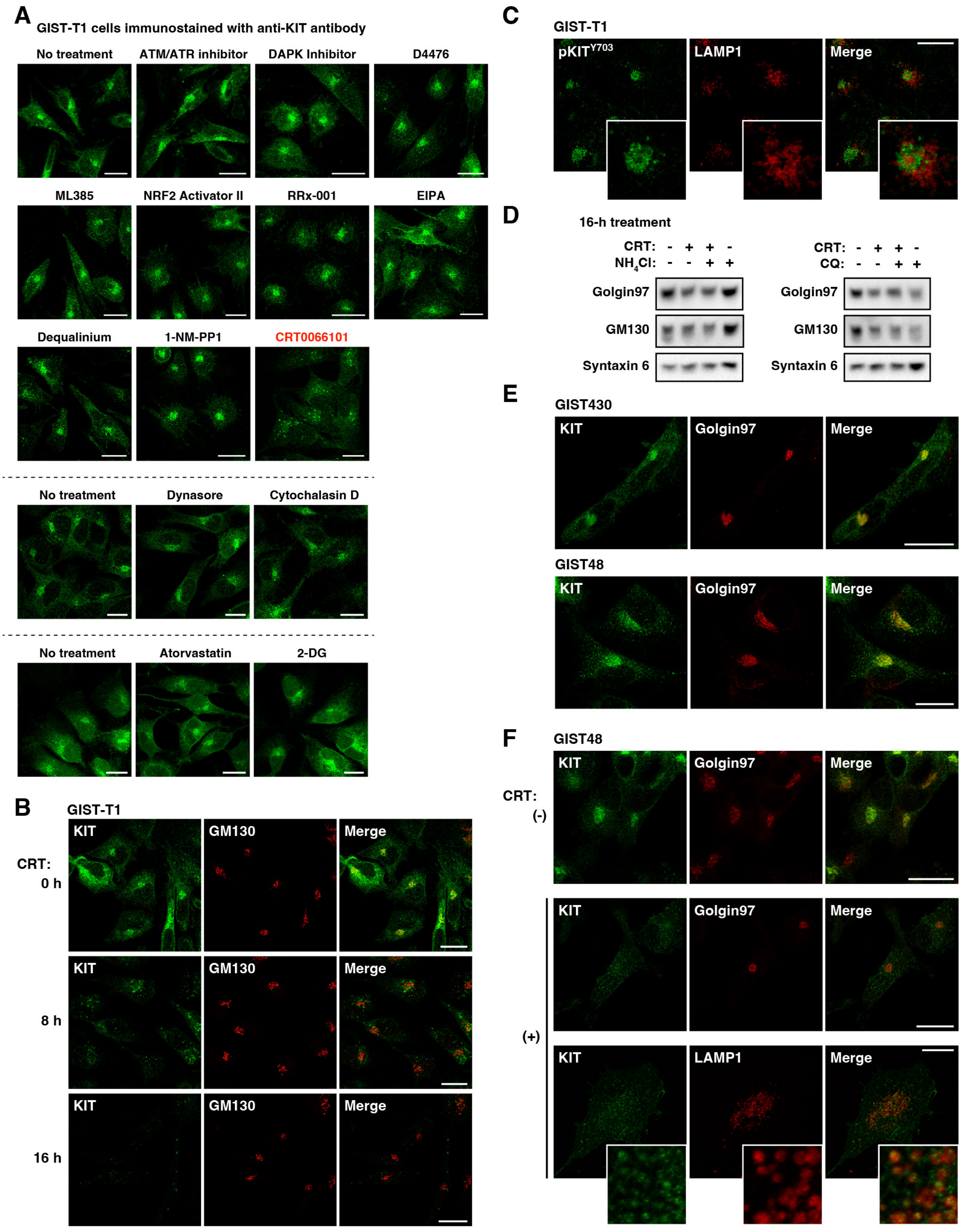


*(legend on next page)*

**Figure S1. KIT requires PKD activity for its Golgi retention in GIST cells, related to Figure 1**

(A) GIST-T1 cells were treated for 8–20 hours with the following inhibitors: 10 μM ATM/ATR kinase inhibitor, 100 μM DAPK inhibitor, 100 μM D4476 (casein kinase I inhibitor), 100 μM ML385 (NRF2 inhibitor), 100 μM NRF2 activator II, 1 μM RRX-001 (G6PD inhibitor), 20 μM EIPA (NHE inhibitor), 5 μM dequalinium chloride (PKC inhibitor), 40 μM 1-NM-PP1 (SRC inhibitor), 10 μM CRT0066101 (PKD inhibitor), 100 μM Dynasore (dynamin inhibitor) or 500 nM cytochalasin D (inhibitor of actin polymerization), 100 μM atorvastatin (HMG-CoA reductase inhibitor), or 10 mM 2-DG (glycolysis inhibitor). Cells were immunostained for KIT. NB: PKD inhibitor CRT0066101 decreased perinuclear KIT. Bars, 20 μm.

(B) GIST-T1 cells were treated with 10 μM CRT0066101 (CRT) for the indicated periods, then immunostained for KIT (green) and GM130 (Golgi matrix protein 130 kDa, red). Bars, 20 μm.

(C) GIST-T1 cells were immunostained for phospho-$KIT^{Y703}$ ($pKIT^{Y703}$, green) and LAMP1 (lysosomal marker, red). Magnified images of the perinuclear region are shown. Bar, 20 μm.

(D) GIST-T1 cells were treated with 10 μM CRT and/or 20 mM $NH_4Cl$, 100 μM chloroquine (CQ) for 16 hours, then immunoblotted for golgin97, GM130, and syntaxin 6. Total protein levels were confirmed by Coomassie Brilliant Blue staining as in the main Figure 1F.

(E) GIST430 or GIST48 cells were immunostained with anti-KIT (green) and anti-golgin97 (red). Bars, 20 μm.

(F) GIST48 cells were treated with 20 μM CRT for 16 hours, then immunostained for KIT (green) in conjunction with golgin97 (red) or LAMP1 (red). Magnified images of the lysosomal region are shown. Bars, 20 μm.

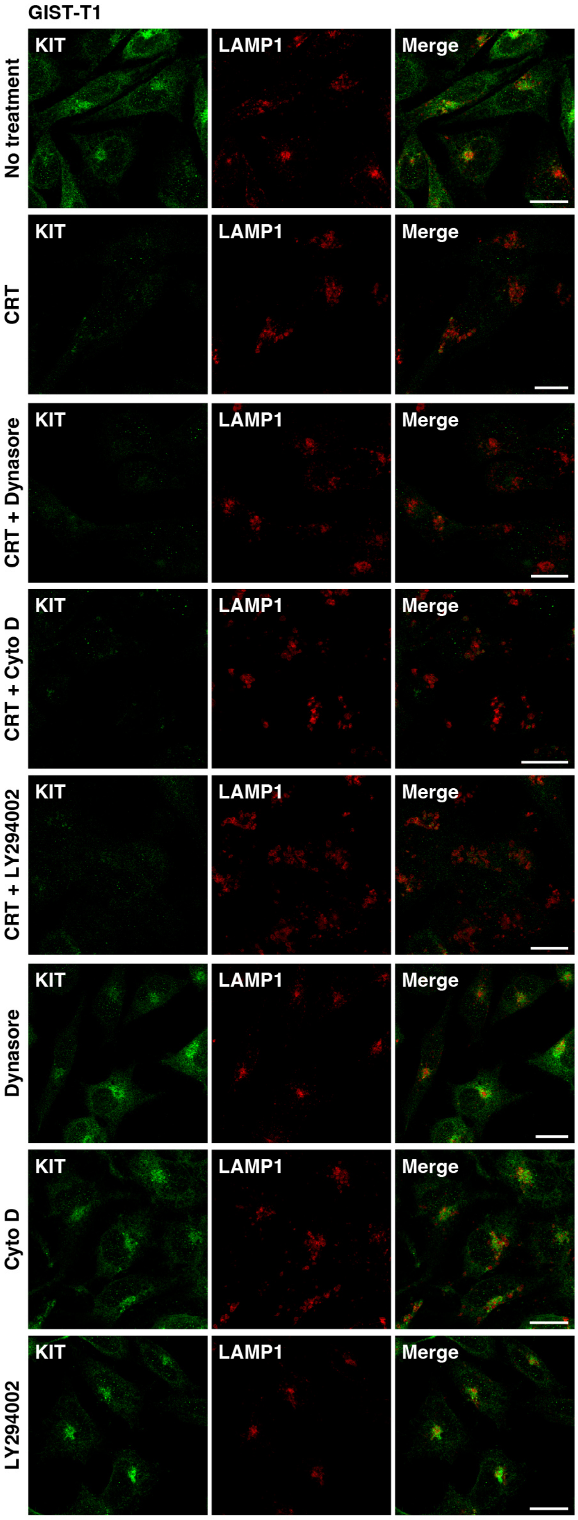


**Figure S2. Dynasore, cytochalasin D, and LY294002 have no effect on KIT trafficking in GIST-T1 cells, related to Figure 2**
GIST-T1 cells were treated with 10 $\mu$M CRT0066101 (CRT, PKD inhibitor) and/or 100 $\mu$M dynasore (dynamin inhibitor), 500 nM cytochalasin D (inhibitor of actin polymerization), and 20 $\mu$M LY294002 (PI3-kinase inhibitor) for 16 hours. Cells were immunostained for KIT (green) and LAMP1 (lysosomal marker, red). Bars, 20 $\mu$m.

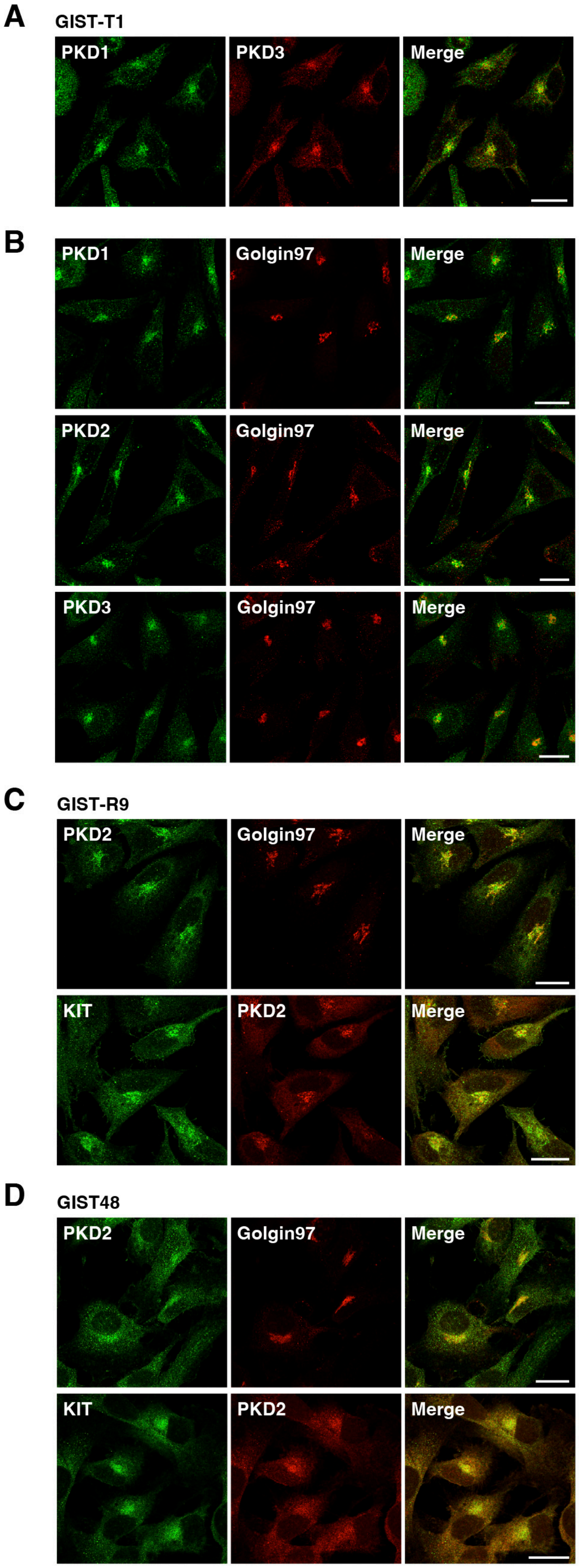


**Figure S3. PKD2 localizes to the Golgi area in GIST cells, related to Figure 3**

(A and B) GIST-T1 cells were immunostained for golgin97 (red) in conjunction with PKD1 (green), PKD2 (green), or PKD3 (red or green). Bars, 20 $\mu$m.

(C and D) GIST-R9 (C) and GIST48 cells (D) were immunostained for PKD2 (green or red) in conjunction with golgin97 (red) or KIT (green). Bars, 20 $\mu$m.

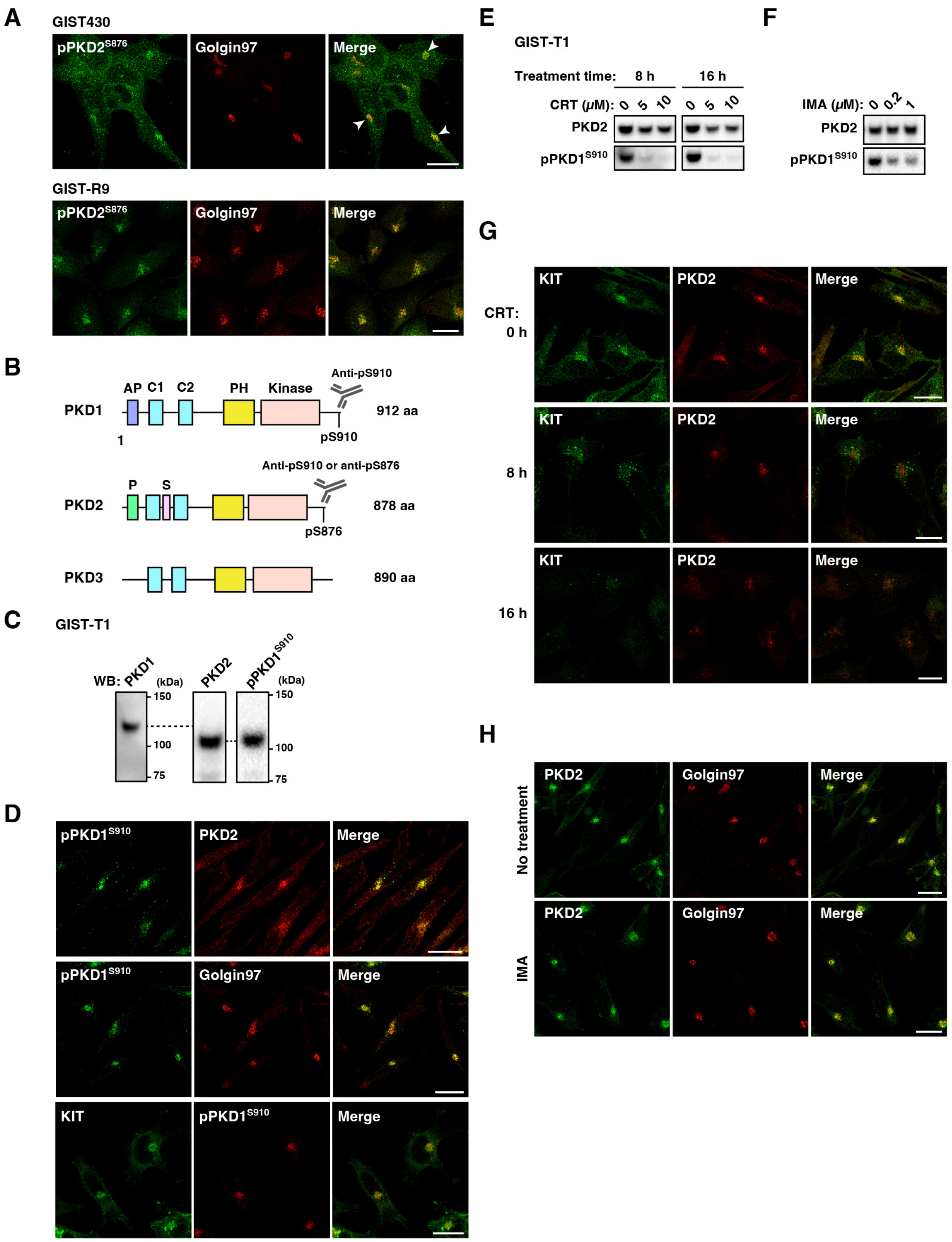


*(legend on next page)*

**Figure S4. PKD2 is autophosphorylated in the Golgi area in GIST cells, related to Figure 4**

(A) GIST430 and GIST-R9 cells were immunostained for phospho-PKD2$^{S876}$ (pPKD2$^{S876}$, green) and golgin97 (red). Arrowheads indicate the Golgi region in GIST430 cells. Bars, 20 $\mu$m.

(B) Schematic representations of PKD members showing the C1a, C1b domains (diacylglycerol-binding domains), the pleckstrin homology (PH) domain, and the kinase domain. AP, alanine- and proline-rich domain; P, proline-rich domain; S, serine-rich domain. NB: Anti-pPKD1$^{S910}$ binds to both pS910 in PKD1 and pS876 in PKD2.

(C) GIST-T1 lysates were immunoblotted with anti-PKD1, anti-PKD2, and anti-pPKD1$^{S910}$ antibodies. NB: In GIST-T1, anti-pPKD1$^{S910}$ mainly reacted with PKD2.

(D) GIST-T1 cells were immunostained with anti-pPKD1$^{S910}$ (green or red) in conjunction with PKD2 (red), golgin97 (red), or KIT (green). Bars, 20 $\mu$m.

(E) GIST-T1 cells were treated with CRT0066101 (CRT, PKD inhibitor) for the indicated periods, and immunoblotted.

(F) GIST-T1 cells were treated with 200 nM imatinib (IMA, KIT kinase inhibitor) for 8 hours, then immunoblotted with anti-PKD2 and anti-pPKD1$^{S910}$ antibodies.

(G) GIST-T1 cells were treated with 10 $\mu$M CRT for the indicated periods. Cells were immunostained for PKD2 (green) and golgin97 (red). Bars, 20 $\mu$m. NB: PKD2 was found in the Golgi area in the presence of CRT.

(H) GIST-T1 cells were treated with 200 nM IMA for 8 hours, then immunostained for PKD2 (green) and golgin97 (red). Bars, 20 $\mu$m.

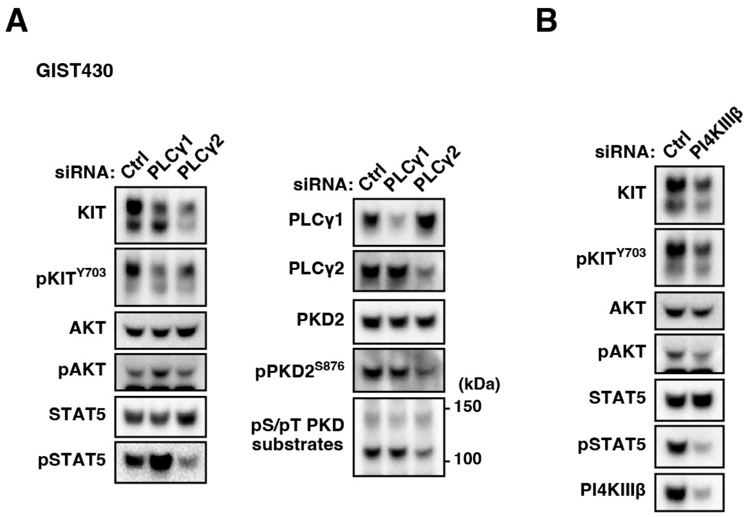


**Figure S5. Knockdown of PLCγ2/PI4KIIIβ also reduces the protein levels of KIT in GIST430 cells, related to Figure 5 and Figure 6**
(A and B) GIST430 cells were transfected with siRNA targeting (A) PLCγ1, PLCγ2, or (B) PI4KIIIβ for 30 hours, and immunoblotted.

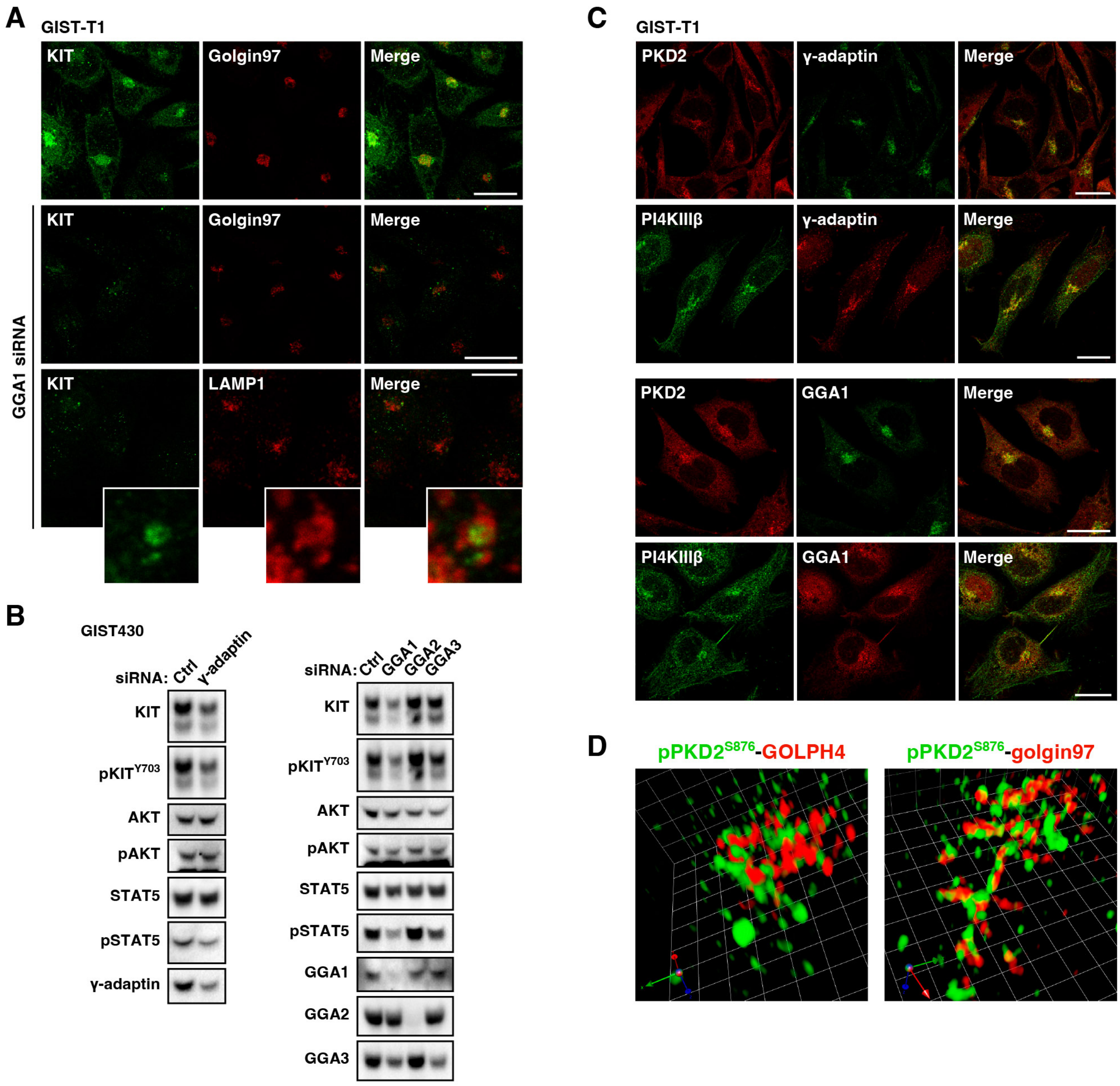


**Figure S6. AP1 and GGA1 play a critical role in KIT retention in GIST cells, related to Figure 7**

(A) GIST-T1 cells were transfected with the indicated siRNAs for 30 hours. Cells were immunostained for KIT (green) in conjunction with golgin97 (red) or LAMP1 (lysosomal marker, red). Magnified images of lysosomal region are shown. Bars, 20 $\mu$m.

(B) GIST430 cells were transfected with the indicated siRNAs for 30 hours, then immunoblotted.

(C) GIST-T1 cells were immunostained with the indicated antibodies. Bars, 20 $\mu$m.

(D) GIST-T1 cells were immunostained for phospho-PKD2$^{S876}$ (pPKD2$^{S876}$, green) in conjunction with Golgi phosphoprotein 4 (GOLPH4, *medial*-Golgi marker, red) or golgin97 (*trans*-Golgi network marker, red). Stained proteins were visualized with the super-resolution confocal live imaging microscopy (SCLIM) developed in RIKEN. Representative 3D images are shown. On side of the grid in the right and left panels indicates 1.49 $\mu$m and 1.58 $\mu$m, respectively.

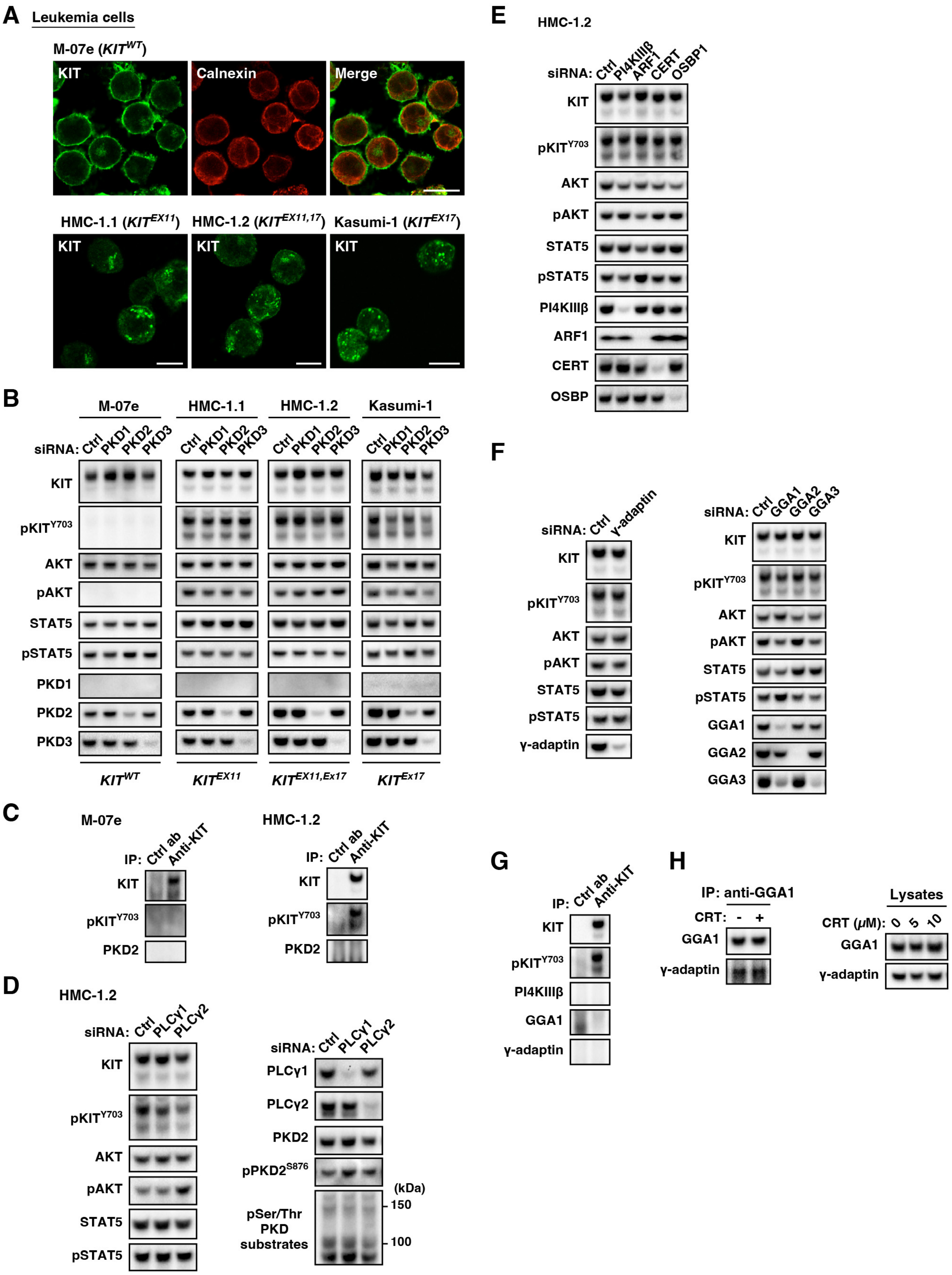


**Figure S7. PKD2 does not play a role in the intracellular retention of KIT and the mutant signaling in leukemia cells**

(A) M-07e, HMC-1.1, HMC-1.2, and Kasumi-1 cells were immunostained with the indicated antibodies. Bars, 10 $\mu$m.

(B) Leukemia cells were transfected with the indicated siRNAs for 48 hours, and immunoblotted.

(C) KIT was immunoprecipitated from M-07e (left) or HMC-1.2 cells (right). Immunoprecipitates (IPs) were immunoblotted.

(D–F) HMC-1.2 cells were transfected with the indicated siRNAs, and immunoblotted.

(G) KIT in HMC-1.2 was immunoprecipitated with anti-KIT antibody. IPs were immunoblotted.

(H) HMC-1.2 cells were treated with 10 $\mu$M CRT0066101 (CRT, PKD inhibitor) for 8 hours. GGA1 was immunoprecipitated. IPs were immunoblotted.

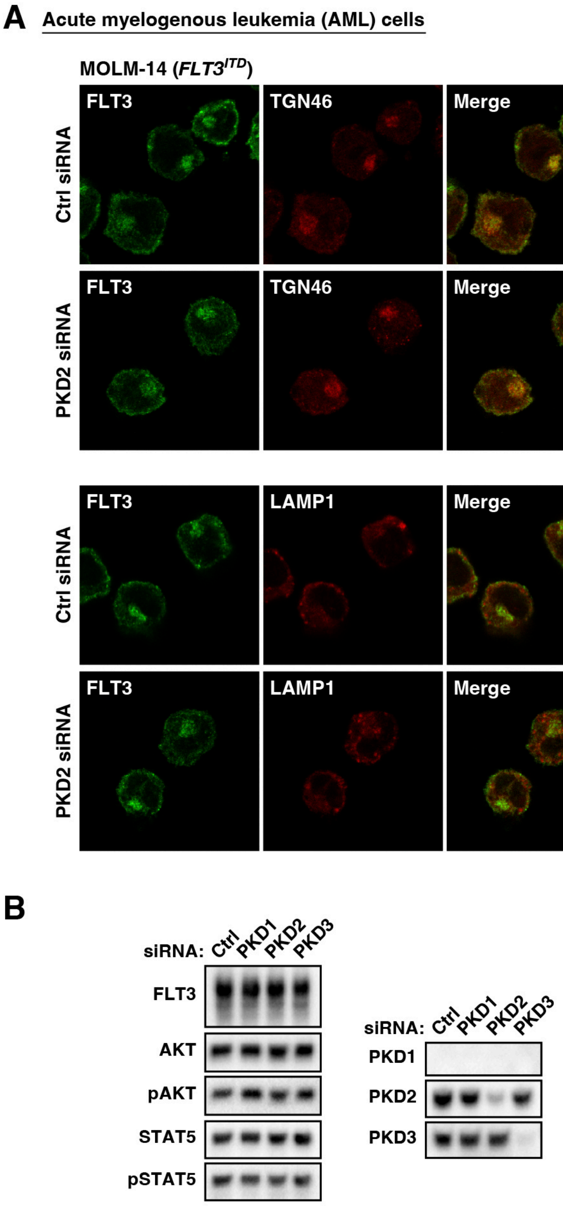


**Figure S8. FLT3-ITD is retained in the Golgi area in a PKD-independent manner in AML cells**

(A and B) MOLM-14 cells were transfected with the indicated siRNAs for 48 hours. (A) Cells were immunostained for FLT3 (green) in conjunction with TGN46 (*trans*-Golgi network marker, red) or LAMP1 (lysosomal marker, red). Bars, 10 $\mu$m. (B) Lysates were immunoblotted.

| Antibody | Host | Clone/catalog # | Distribution source |
|---|---|---|---|
| AKT | Mouse | 40D4 | Cell Signaling Technology |
| AKT [pT308] | Rabbit | C31E5E | Cell Signaling Technology |
| ARF1 | Mouse | ARFS 1A9/5 | Santa Cruz Biotechnology |
| Calnexin | Rabbit | ADI-SPA-860 | Enzo |
| CERT | Rabbit | 15191-1-AP | Proteintech |
| FLT3 | Rabbit | S-18 | Santa Cruz Biotechnology |
| FLT3 | Mouse | SF1.340 | Santa Cruz Biotechnology |
| γ-adaptin | Mouse | F-10 | Santa Cruz Biotechnology |
| γ-adaptin | Mouse | 88 | BD Transduction Laboratories |
| γ-adaptin | Rabbit | 13258-1-AP | Proteintech |
| GGA1 | Rabbit | 25674-1-AP | Proteintech |
| GGA1 | Mouse | D-6 | Santa Cruz Biotechnology |
| GGA2 | Mouse | E-3 | Santa Cruz Biotechnology |
| GGA2 | Mouse | 27 | BD Transduction Laboratories |
| GGA3 | Mouse | 8 | Santa Cruz Biotechnology |
| GGA3 | Mouse | 8 | BD Transduction Laboratories |
| GM130 | Mouse | 35 | BD Transduction Laboratories |
| Golgin97 | Rabbit | D8P2K | Cell Signaling Technology |
| Golgin97 | Mouse | CDF4 | Thermo Fisher Scientific |
| GOLPH4 | Mouse | XY-2 | Santa Cruz Biotechnology |
| KIT | Rabbit | D13A2 | Cell Signaling Technology |
| KIT | Rabbit | D3W6Y | Cell Signaling Technology |
| KIT | Mouse | ab81 | Cell Signaling Technology |
| KIT | Goat | M-14 | Santa Cruz Biotechnology |
| KIT | Mouse | E-1 | Santa Cruz Biotechnology |
| KIT | Mouse | E-3 | Santa Cruz Biotechnology |
| KIT | Mouse | 104D2 | BioLegend |
| KIT | Mouse | 28 | BD Transduction Laboratories |
| KIT [pY703] | Rabbit | D12E12 | Cell Signaling Technology |
| LAMP1 | Mouse | D4O1S | Cell Signaling Technology |
| LAMP1 | Rabbit | L1418 | Sigma-Aldrich |

**Supplementary Table 1. List of primary antibodies.**

| Antibody | Host | Clone/catalog # | Distribution source |
|---|---|---|---|
| OSBP1 | Mouse | A-5 | Santa Cruz Biotechnology |
| PDGFRA | Rabbit | D13C6 | Cell Signaling Technology |
| PI4KIIIβ | Mouse | E-4 | Santa Cruz Biotechnology |
| PI4KIIIβ | Mouse | 7 | BD Transduction Laboratories |
| PI4KIIIβ | Rabbit | 13247-1-AP | Proteintech |
| PI4P | Mouse | Z-P004 | Echelon Biosciences |
| PKD1 | Mouse | ab172096 | Abcam |
| PKD1 | Rabbit | D4J1N | Cell Signaling Technology |
| PKD1 [pS916] | Rabbit | #2051 | Cell Signaling Technology |
| PKD2 | Rabbit | D1A7 | Cell Signaling Technology |
| PKD2 | Mouse | F-2 | Santa Cruz Biotechnology |
| PKD2 | Mouse | O95G1 | BioLegend |
| PKD2 [pS876] | Rabbit | EP1496Y | Abcam |
| PKD2 [pS876] | Rabbit | SAB4504104 | Sigma-Aldrich |
| PKD3 | Rabbit | D57E6 | Cell Signaling Technology |
| PLCγ1 | Mouse | E-12 | Santa Cruz Biotechnology |
| PLCγ2 | Rabbit | E5U4T | Cell Signaling Technology |
| PLCγ2 | Mouse | 2F8H5 | Proteintech |
| PLCγ2 [pY759] | Rabbit | #3874 | Cell Signaling Technology |
| PLCγ2 [pY759] | Rabbit | E9E9Y | Cell Signaling Technology |
| pS/pT PKD substrate | Rabbit | #4381 | Cell Signaling Technology |
| pTyr | Rabbit | pY1000 | Cell Signaling Technology |
| pTyr | Mouse | 4G10 | Millipore |
| STAT5 | Rabbit | C-17 | Santa Cruz Biotechnology |
| STAT5 | Rabbit | D2O6Y | Cell Signaling Technology |
| STAT5 | Mouse | 89 | BD Transduction Laboratories |
| STAT5 [pY694] | Rabbit | D47E7 | Cell Signaling Technology |
| Syntaxin 6 | Mouse | 30 | BD Transduction Laboratories |
| TGN46 | Rabbit | ab76282 | Abcam |

**Supplementary Table 2. List of primary antibodies.**

| Antibody | Host | Clone/catalog # | Distribution source |
|---|---|---|---|
| HRP anti-mouse IgG | Donkey | 715-035-151 | Jackson Immuno Research |
| HRP anti-rabbit IgG | Donkey | 711-035-152 | Jackson Immuno Research |
| HRP anti-goat IgG | Donkey | 705-035-147 | Jackson Immuno Research |
| AF488 anti-mouse IgG | Donkey | A21202 | Thermo Fisher Scientific |
| AF488 anti-rabbit IgG | Donkey | A21206 | Thermo Fisher Scientific |
| AF488 Plus anti-rabbit IgG | Donkey | A32790 | Thermo Fisher Scientific |
| AF488 anti-goat IgG | Donkey | A21447 | Thermo Fisher Scientific |
| AF568 anti-mouse IgG | Donkey | A10037 | Thermo Fisher Scientific |
| AF555 Plus anti-mouse IgG | Donkey | A32773 | Thermo Fisher Scientific |
| AF568 anti-rabbit IgG | Donkey | A10042 | Thermo Fisher Scientific |
| AF647 anti-mouse IgG | Donkey | A31571 | Thermo Fisher Scientific |
| AF647 anti-rabbit IgG | Donkey | A31573 | Thermo Fisher Scientific |
| AF488 anti-mouse IgM | Goat | A21042 | Thermo Fisher Scientific |
| AF647 lectin-HPA | - | L32454 | Thermo Fisher Scientific |

**Supplementary Table 3. List of secondary antibodies.** HRP, horseradish peroxidase; AF, Alexa Fluor.

| Compound | Catalog # | Solvent | Distribution source |
|---|---|---|---|
| 1-NM-PP1 | HY-13942 | DMSO | MedChemExpress |
| 2-deoxy-D-glucose | D6134 | Water | Sigma-Aldrich |
| Ammonium chloride | A9434 | Water | Sigma-Aldrich |
| ATM/ATR inhibitor | 118501 | DMSO | Sigma-Aldrich |
| Atorvastatin calcium salt trihydrate | PZ0001 | DMSO | Sigma-Aldrich |
| Chloroquine diphosphate salt | C6628 | Water | Sigma-Aldrich |
| CRT0066101 dihydrochloride | S8366 | DMSO | Selleck |
| Cytochalasin D | C8273 | DMSO | Sigma-Aldrich |
| D4476 | S7642 | DMSO | Selleck |
| DAPK Inhibitor (TC-DAPK 6) | 324788 | DMSO | Sigma-Aldrich |
| Dequalinium chloride | 048-30591 | Water | Wako Chemicals |
| Dynasore hydrate | D7693 | DMSO | Sigma-Aldrich |
| EIPA | HY-101840 | DMSO | MedChemExpress |
| Imatinib mesylate | 13139 | DMSO | Cayman Chemical |
| PI4KIIIβ-IN-10 | HY-100198 | DMSO | MedChemExpress |
| LY294002 | 440202 | DMSO | Sigma-Aldrich |
| ML385 | HY-100523 | DMSO | MedChemExpress |
| NRF2 Activator II | 492041 | DMSO | Sigma-Aldrich |
| PIK-93 | S1489 | DMSO | Selleck |
| RRx-001 | S8405 | DMSO | Selleck |

**Supplementary Table 4. List of inhibitors and activators.**

| Target | Catalog # | Distribution source |
|---|---|---|
| PKD1 | L-005028-00-0005 | Dharmacon |
| PKD2 | L-004197-00-0005 | Dharmacon |
| PKD3 | L-005029-00-0005 | Dharmacon |
| PLCγ1 | L-003559-00-0005 | Dharmacon |
| PLCγ2 | L-008339-02-0005 | Dharmacon |
| PI4KIIIβ | L-006777-00-0005 | Dharmacon |
| ARF1 | L-011580-00-0005 | Dharmacon |
| CERT | L-012101-00-0005 | Dharmacon |
| OSBP1 | L-009747-00-0005 | Dharmacon |
| γ-adaptin | L-019183-00-0005 | Dharmacon |
| GGA1 | L-013694-00-0005 | Dharmacon |
| GGA2 | L-012908-01-0005 | Dharmacon |
| GGA3 | L-012881-01-0005 | Dharmacon |
| Non-targeting control siRNAs | D-001810-10-20 | Dharmacon |

**Supplementary Table 5. List of siRNAs.**
